# Predicting master transcription factors from pan-cancer expression data

**DOI:** 10.1101/839142

**Authors:** Jessica Reddy, Marcos A. S. Fonseca, Rosario I. Corona, Robbin Nameki, Felipe Segato Dezem, Isaac A. Klein, Heidi Chang, Daniele Chaves-Moreira, Lena Afeyan, Tathiane M Malta, Xianzhi Lin, Forough Abbasi, Alba Font-Tello, Thais Sabedot, Paloma Cejas, Norma Rodríguez-Malavé, Ji-Heui Seo, De-Chen Lin, Ursula Matulonis, Beth Y. Karlan, Simon A. Gayther, Alexander Gusev, Houtan Noushmehr, Henry Long, Matthew L. Freedman, Ronny Drapkin, Brian J. Abraham, Richard A. Young, Kate Lawrenson

## Abstract

The function of critical developmental regulators can be subverted by cancer cells to control expression of oncogenic transcriptional programs. These “master transcription factors” (MTFs) are often essential for cancer cell survival and represent vulnerabilities that can be exploited therapeutically. The current approaches to identify candidate MTFs examine super-enhancer associated transcription factor-encoding genes with high connectivity in network models. This relies on chromatin immunoprecipitation-sequencing (ChIP-seq) data, which is technically challenging to obtain from primary tumors, and is currently unavailable for many cancer types and clinically relevant subtypes. In contrast, gene expression data are more widely available, especially for rare tumors and subtypes where MTFs have yet to be discovered. We have developed a predictive algorithm called CaCTS (Cancer Core Transcription factor Specificity) to identify candidate MTFs using pan-cancer RNA-sequencing data from The Cancer Genome Atlas. The algorithm identified 273 candidate MTFs across 34 tumor types and recovered known tumor MTFs. We also made novel predictions, including for cancer types and subtypes for which MTFs have not yet been characterized. Clustering based on MTF predictions reproduced anatomic groupings of tumors that share 1-2 lineage-specific candidates, but also dictated functional groupings, such as a squamous group that comprised five tumor subtypes sharing 3 common MTFs. *PAX8, SOX17*, and *MECOM* were candidate factors in high-grade serous ovarian cancer (HGSOC), an aggressive tumor type where the core regulatory circuit is currently uncharacterized. *PAX8, SOX17*, and *MECOM* are required for cell viability and lie proximal to super-enhancers in HGSOC cells. ChIP-seq revealed that these factors co-occupy HGSOC regulatory elements globally and co-bind at critical gene loci including *MUC16* (CA-125). Addiction to these factors was confirmed in studies using THZ1 to inhibit transcription in HGSOC cells, suggesting early down-regulation of these genes may be responsible for cytotoxic effects of THZ1 on HGSOC models. Identification of MTFs across 34 tumor types and 140 subtypes, especially for those with limited understanding of transcriptional drivers paves the way to therapeutic targeting of MTFs in a broad spectrum of cancers.

## Introduction

Accumulating evidence indicates that tumor cells are driven by a small set of transcription factors (TFs) that control global gene expression programs (Durbin et al., 2018; Mansour et al., 2014; Sanda et al., 2012). While disused in corresponding healthy cells, some tumor-driving master transcription factors (MTFs) are often developmental regulators that are aberrantly expressed and functionally co-opted to regulate tumor cell states. For example, regulators of T cell development, TAL1, GATA3, RUNX1, and MYB, are over-expressed and co-regulate oncogenic programs in T-cell acute lymphoblastic leukemias (Mansour et al., 2014; Sanda et al., 2012). Additionally, developmental regulators MYCN, HAND2, ISL1, PHOX2B, GATA3, and TBX2 have been identified as MTFs in neuroblastoma (Durbin et al., 2018). MTFs are generally expressed in only a limited number of cell types, consistent with their potent role in establishing a gene expression program that drives cell identity (D’Alessio et al., 2015; Whyte et al., 2013). MTFs are a class of promising therapeutic targets as they are selective essentialities in cancer cells, due to a phenomenon termed transcriptional oncogene addiction (Bradner et al., 2017).

Although TFs are notoriously difficult to directly target with small molecules, several MTFs have been shown to be highly sensitive to pharmaceutical inhibition of general transcriptional regulators, including those that target bromodomain (BRD)-containing proteins and transcriptional cyclin-dependent kinase 7 (Chapuy et al., 2013; Durbin et al., 2018; Kwiatkowski et al., 2014; Wang et al., 2015). The expression of tumor cell MTFs are often driven by large clusters of enhancers, or termed super- or stretch-enhancers (SEs) (Lovén et al., 2013; Parker et al., 2013). The exquisite sensitivity of these factors to chemical perturbation of BRDs and transcriptional CDKs is hypothesized to result from disruption of continuous, high-level transcription at super-enhancers (SEs), combined with the short transcript half-lives and auto-regulatory activities. Together, studies on MTFs and transcriptional inhibition in tumor cells demonstrate the importance of identifying these critical factors and investigating if they can be indirectly targeted with small molecules targeting general regulators of transcription. MTFs are thought to form core transcriptional regulatory circuitries by co-occupying genomic sites, particularly at SEs, and co-regulating the expression of MTF genes and others genes critical for cellular identity (Chen et al., 2019a; Durbin et al., 2018; Sanda et al., 2012). Presently, the main approaches to identifying MTFs in cancer cells attempts to model these features by identifying TFs predicted to exhibit evidence of interconnected autoregulation (Ott et al., 2018; Saint-André et al., 2016; Zhang et al., 2018). This involves performing ChIP-seq experiments to map enhancers and identifying SE associated TFs whose predicted binding motifs are enriched at SEs, both upstream and regulating other MTFs (Federation et al., 2018). These approaches have been shown to recover experimentally validated MTFs in various tumor types (Chen et al., 2019a, 2019b; Lin et al., 2016; Ott et al., 2018).

For many tumor tissues, obtaining adequate amounts of primary tumor cells for ChIP-seq experiments can be technically challenging. RNA-sequencing (RNA-seq) experiments, however, require less starting material and, RNA-seq data from primary tumor samples are currently more widely available, especially for rare tumor types and subtypes. We therefore developed an approach to predict tumor MTFs in numerous cancer types and subtypes using RNA-seq data from The Cancer Genome Atlas (TCGA). This approach is called the Cancer Core Transcription factor Specificity (CaCTS) algorithm, and it attempts to determine critical TFs by identifying those exhibiting tumor-specific expression compared to a background data set that contains a diverse set of cancer types. A similar approach has been previously applied to normal tissues (D’Alessio et al., 2015). We find that candidate MTFs identified through the CaCTS algorithm possess many expected qualities of MTFs, such as SE association and high levels of essentiality, indicating our approach is an orthogonal metric to existing attempts to predict MTFs. This unique resource represents a collection of candidate MTFs for 34 tumor types and 140 molecular and histologic subtypes that can be directly explored for therapeutic potential.

## Results

### The Cancer Core Transcription factor Specificity (CaCTS) algorithm identifies factors with features attributed to tumor cell MTFs

Given that many tumor cell MTFs are developmental regulators that exhibit cell-type-specific expression (Durbin et al., 2018; Mansour et al., 2014; Sanda et al., 2012), we hypothesized that these MTFs could be retrieved by identifying sets of TFs that exhibit tumor-specific RNA expression within a diverse collection of tumor types. We therefore mined TCGA RNA-sequencing data containing 9,691 patient samples representing 34 tumor types. To identify candidate MTFs, we developed a method, the Cancer Core Transcription Factor Specificity (CaCTS) algorithm, to identify factors over-expressed in each tumor type relative to the other tumor types represented in the TCGA data set (Figure 1A, Table S1). This approach uses an entropy-based measure of Jensen-Shannon divergence similar to that used to identify candidate MTFs in normal human tissues (D’Alessio et al., 2015). We calculated the average expression levels of 1,578 TFs in 34 tumor types/major subtypes (Table S2). On average, 309 samples (range: 45-1,083) were used to calculate average TF expression values in each of the 34 tumor types. The specificity of expression of each TF, or ‘CaCTS score’, was calculated by comparing its expression level in the query tumor type to that in the remaining 33 tumor types. A high CaCTS score is therefore assigned to factors with high level expression in the query tumor types as compared to background data set (for example, TF_1_ depicted in Figure 1A compared to TF_2_, which represents a factor that is ubiquitously expressed across the cohort). The output of the CaCTS algorithm is a list of all TFs ranked by CaCTS scores in each of the 34 tumor types (Table S3).

**Figure 1.**
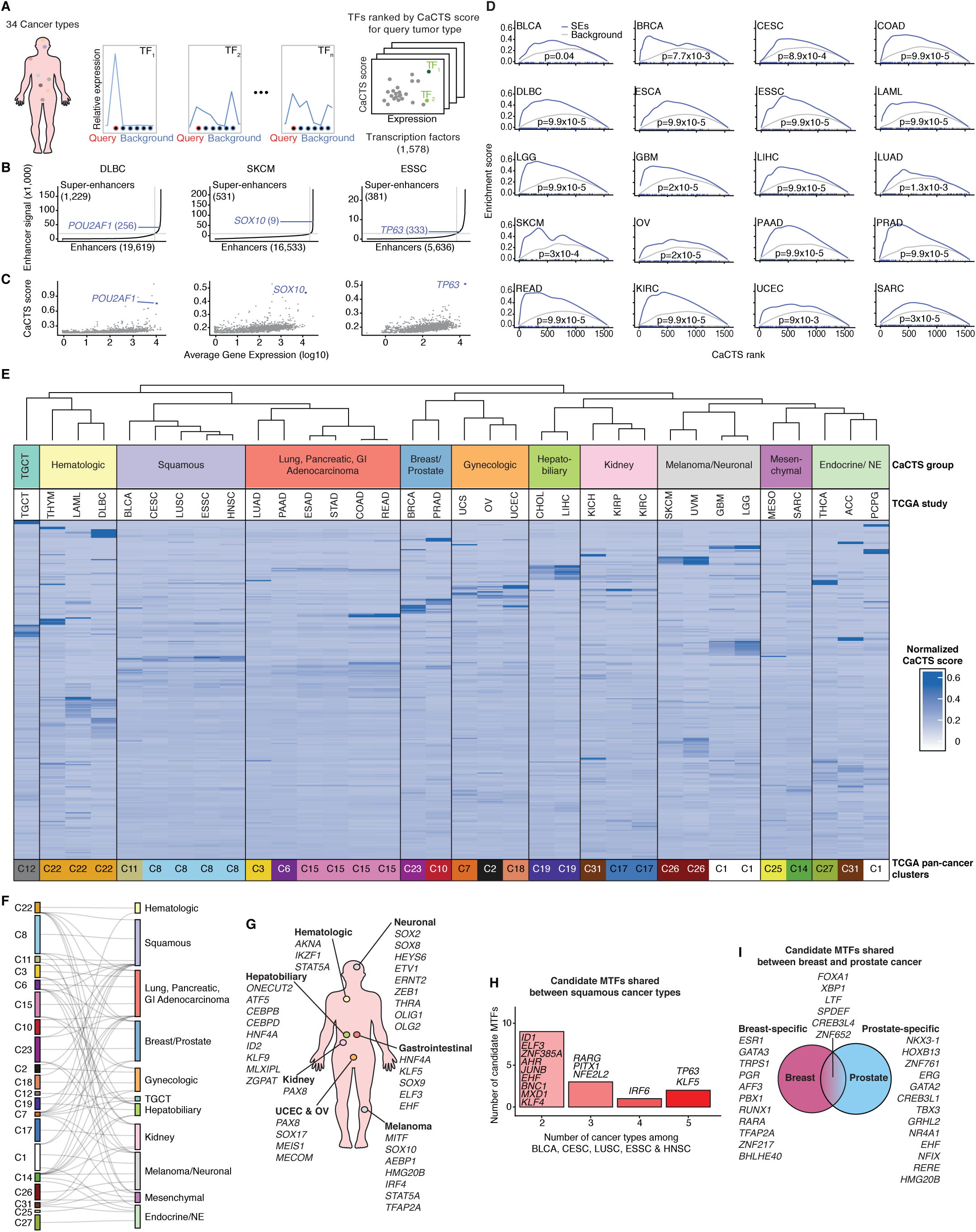
A multi-cancer compendium of candidate master transcription factors. (A) Schematic of the Cancer Core Transcription factor Specificity (CaCTS) approach. (B) Positive control factors - *POU2AF1* in DLBC, *SOX10* in SKCM and *TP63* in ESSC - coincide with large super-enhancers in the relevant tumor tissues. (C) These factors are highly expressed and have large CaCTS scores. (D) Across 20 tumor types, candidate MTFs with high CaCTS scores are significantly enriched at super-enhancers in the corresponding tumor types. Gene-set enrichment p-values are shown, analyses were performed with 10,000 permutations of randomly selected factors. (E) Unsupervised hierarchical clustering based on candidate MTF CaCTS scores across 34 tumor types, using the *spearman* method and *complete* distance parameters. A height cut-off equal to 0.63 defines 11 CaCTS clusters. (F) Sankey plot showing pan-cancer clusters (Hoadley et al., 2018) that correspond to the 11 CaCTS clusters. We identified the TCGA cluster assigned in 50% or more of the tumors in each of our 11 clusters. (G) Factors are often shared across tumor derived from organ systems with a shared development lineage. (H) Squamous tumors from diverse organ sites share keratinocyte differentiation TFs as candidates. (I) Breast and prostate adenocarcinomas share six candidate MTFs.

Lineage-specific oncogenic transcription factors often share features attributed to MTFs, including association with SEs (Chapuy et al., 2013; Eliades et al., 2018; Shang et al., 2019; Yuan et al., 2017). Among these include *POU2AF1* (OCA-B) in diffuse large B-cell lymphoma (DLBC) (Chapuy et al., 2013), *SOX10* in melanoma (SKCM) (Eliades et al., 2018), and *TP63* in esophageal squamous carcinoma (ESSC) (Yuan et al., 2017). *POU2AF1* is critical for B-cell fate determination and was found proximal to especially large BRD4-loaded SEs in DLBC cells (Chapuy et al., 2013). *SOX10* is important for melanoma cell proliferation and survival and is associated with SEs in melanoma tissues (Eliades et al., 2018). *TP63* is a known master regulator of keratinocyte differentiation and associated with SEs in esophageal squamous cells (Yuan et al., 2017). By analyzing publicly available H3K27ac data, we confirmed that *POU2AF1, SOX10*, and *TP63* are associated with SEs in DLBC, primary melanoma tissues, and an esophageal squamous cell carcinoma cell line, respectively (Figure 1B, Figure S1A). *POU2AF1, SOX10*, and *TP63* were scored highly with the CaCTS algorithm, ranking fourth, second, and first, in DLBC, SKCM, and ESSC, respectively (Figure 1C, Table S3). Although *TP63* was the top-scoring factor in esophageal squamous tumors, it ranked poorly in esophageal adenocarcinoma (rank 1,392 out of 1,578 TFs), reinforcing previous observations that this factor is a distinguishing feature between the two histologic subtypes (Cancer Genome Atlas Research Network et al., 2017a). These findings demonstrate that the CaCTS algorithm is able to recover factors with known MTF-related roles in tumor cells.

We next sought to investigate if SE associated genes are highly ranked by the CaCTS algorithm in general. We collected publicly available and internal H3K27ac ChIP-seq data sets, collecting an average of 4.5 (range 1-25) samples to represent 20 out of the 34 tumor types (Table S4). Fifteen of these were from primary tumors, while only cell line data were available for BLCA, CESC, LUAD, SARC, ESAD, and ESSC. We were unable to retrieve H3K27ac data sets for ACC, HNSC, KICH, KIRP, LUSC, MESO, PCPG, STAD, TGCT, THCA, THYM, UCS, and UVM, so these tumor types were not included in this analysis. In each data set, we performed peak calling, identified SEs, and performed gene set enrichment analysis to determine if factors with high CaCTS scores tend to be enriched for SE associated TFs. The majority (81/92, 88%) of comparisons between SE and CaCTS ranks showed significant enrichment (p_GSEA_ < 0.05) (Figure 1D, Figure S1B, Table S5). This analysis demonstrates that TFs with high CaCTS ranks tend to be enriched for SE-associated factors in these twenty tumor types and suggests that *bona fide* MTFs are likely present among high CaCTS scoring factors in the 14 tumor types where enhancer data was not available. We noted that most known candidate factors were among the top 5% of expressed factors in the relevant tumor type (Chapuy et al., 2013; Eliades et al., 2018; Yuan et al., 2017). Therefore, to arrive at a collection of candidate MTFs, we retrieved those factors with high CaCTS scores (top 5%; CaCTS rank ≤ 79) that were also highly expressed (within the top 5% of expression; expression rank ≤ 79) for each of the 34 tumor types (Table S6, Figure S1C). This list contained a total of 273 candidate factors with an average of 8.0 candidate factors per tumor type (range 3-31).

### Candidate MTFs are shared among tumors of similar anatomic or functional state

On average, an individual factor was identified as a candidate MTF for 1.9 tumor types (Figure S2A), consistent with our expectation that cancer MTFs will be enriched for lineage-specific developmental regulators. Hierarchical clustering of tumors based on CaCTS scores of the 273 different candidate MTFs identified 11 clusters (Figure 1E). MTFs identified in testicular germ cell tumors (TGCTs) exhibited the highest average CaCTS scores (TGCT average CaCTS score = 0.91±0.73; compared to an average of 0.36±0.25 across the whole cohort). MTF candidates for this tumor type included known TGCT markers *NANOG* (ranked number 1, CaCTS score = 2.4) and *POU51B* (ranked number 5, CaCTS score = 2.4) (Santagata et al., 2007). Most tumors clustered by organ, which was expected because expression of lineage-specific fators should be similar among tumors originating from related sites. Some clusters represented functional categories; tumors with major squamous components clustered together, and tumors arising from hormonal-responsive organs (breast, prostate) and gynecologic tissues formed two closely related clusters. We also identified a cluster consisting of ectoderm-derived adenocarcinomas from the lung, pancreas, esophagus, stomach, colon and rectum (Figure 1E). These functional groupings were less expected, but they suggest shared MTFs across diverse tissue types are responsible for specific cellular functions and differentiation states. Our clusters were largely consistent with those defined by TCGA unsupervised consensus clustering of RNA-seq data from 10,165 tumor samples (Hoadley et al., 2018) (Figure 1F; Figure S2B). Our set of 273 candidates could successfully recapitulate clusters derived from analysis of ∼15,000 genes, suggesting our predictions contain key drivers of global gene expression programs. There were some notable differences between TCGA pan-cancer clusters and our own. In our clustering, urothelial bladder carcinoma (BLCA) clusters with squamous tumors, while lung and pancreatic adenocarcinomas now cluster with other gastrointestinal solid tumors. This argues that common factors or related factors may manifest in different expression programs depending on cellular context.

Clusters are driven by CaCTS score and so both candidates and non-candidates will influence the clustering; but we identified a subset of 62 MTFs that were shared among 3 or more tumor types (Figure S2A) and sought to further examine commonalities across the union set of candidates (Figure 1E,G). The most striking functional group is squamous, where 3 factors - *TP63, IRF6, KLF5* - were common candidates among 5 tumors from diverse anatomic sites - bladder, cervix, lung, esophagus and head and neck (Figure 1E,H). Six factors were shared between breast and prostate cancer, and both are derived from hormonal responsive organs (*FOXA1, XBP1, LTF, SPDEF, CREB3L4* and *ZNF652*) (Figure 1I). *FOXA1* is a critical player in breast and prostate cancer risk and somatic development (Ciriello et al., 2015; Grasso et al., 2012; Pomerantz et al., 2015; Robinson and Carroll, 2012); *XBP1* is involved in the unfolded protein response and is part of the MYC signaling axis in both cancer types (Zhao et al., 2018); *SPDEF*, also known as prostate-derived Ets factor (*PDEF*) may function as a tumor suppressor gene in prostate cancer, but as an oncogene in breast cancer, where it regulators expression of lineage-specific genes in mammary luminal epithelial cells (Buchwalter et al., 2013; Cheng et al., 2014). *LTF, CREB3L4* and *ZNF652* have been less well studied and warrant further investigation as BRCA and PRAD candidate MTFs. While breast and prostate cancer closely clustered with gynecologic tumors arising from other hormonal-responsive organs, these tumor types shared more candidate factors with each other than with ovarian serous cystadenocarcinoma (OV), uterine carcinosarcoma (USC) and uterine corpus endometrial carcinoma (UCEC). This is consistent with germline susceptibility studies that report greater pleiotropy between prostate and breast cancer risk than between prostate and ovary (Jiang et al., 2019; Kar et al., 2016).

### Candidate MTFs represent tumor cell dependencies

Tumor cell MTFs are expected to be required for critical cellular processes (Bradner et al., 2017), and they are often required for cellular viability. We next determined the extent to which loss-of-function of our candidate factors affects tumor cell viability by examining CRISPR/Cas9 screening data from The Cancer Dependency Map Consortium (depmap.org) (Meyers et al., 2017; Tsherniak et al., 2017). This resource includes CRISPR knockout screening in representative cell lines for 20 of the 34 tumor types in our study, with a mean of 13 cell lines per tumor type (range 1-31), and 434 cell lines in total. Dependency scores (CERES) were normalized such that −1 corresponds to the median effects of pan-essential genes (Meyers et al., 2017). Dependency scores for the candidate MTFs in relevant cell line models are depicted in Figure 2A-T. We calculated the average dependency scores for positive control MTFs - *POU2AF1* in DLBC, *SOX10* in SKCM, and *TP63* in ESSC; these factors have mean dependency scores of −0.71, −1.05, and −0.36, across 4 DLBC, 30 SKCM, and 19 ESSC cell lines, respectively. On average these 3 factors met −0.4, −0.6, −0.8, and −1.0 gene-dependency score cutoffs in 67%, 55%, 50%, and 28% of cell lines for the relevant tumor type (Figure 2A-C, Figure S3). Given these cutoffs, we calculated the proportion of cell lines for each tumor type that were dependent on CaCTS candidate MTFs. We found that on average 87% of tumor types show at least modest dependency (dependency scores of ≤ −0.4) on at least one candidate MTF in at least 50% of the corresponding cell lines, with 53% of tumor types showing high levels of dependency on one or more candidates in at least 50% of the corresponding cell lines (dependency scores of ≤ −1.0) (Figure 2U). This included novel factors such as *PREB*, a factor that regulates prolactin expression and glucose homeostasis in the liver (Park et al., 2018) and *FOXM1*, previously implicated in ESSC (Song et al., 2018; Takata et al., 2014) although not as a master regulator. Lineage-specific dependency data were available for 9 tumor types (DLBC, SKCM, ESSC, BRCA, COAD, KIRC, LIHC, LUSC and OV); 4-53% of CaCTS candidates were also lineage-specific TF dependencies in the relevant tumor type. This was particularly striking for BRCA, where 9/17 candidates were dependencies specifically enriched in breast cancer (Figure 2E).

**Figure 2.**
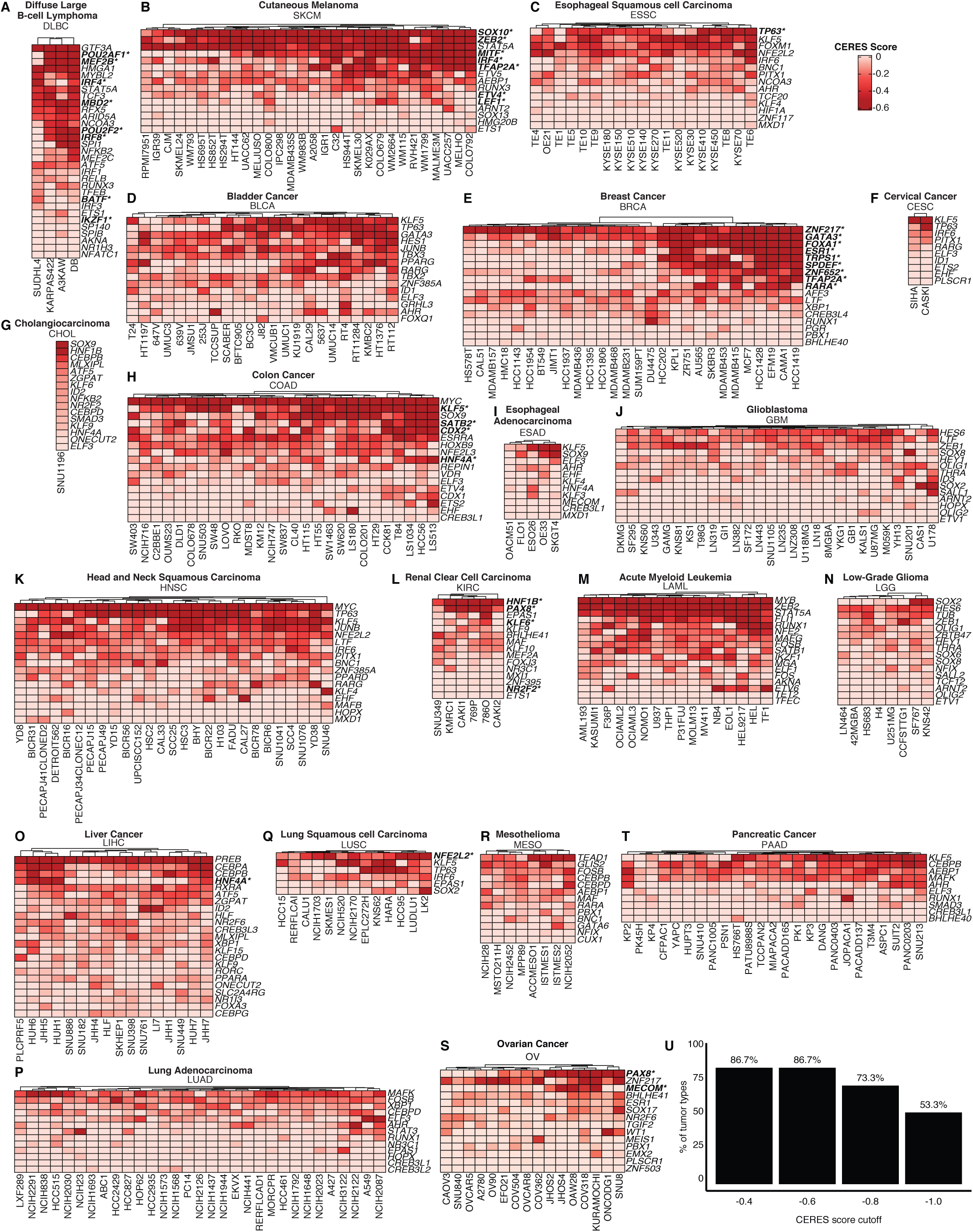
Candidate MTFs are essential genes. Dependency scores for MTF knockouts. Darker red color denotes higher levels of essentiality. Unsupervised hierarchical clustering was used to arrange cell lines (columns) and MTFs (rows). (A)-(C) Tumor types with positive control MTFs indicated: (A) *POU2AF1* in DLBC, (B) *SOX10* in SKCM, and (C) *TP63* in ESSC. (D)-(T) Dependencies for MTF candidates across 17 tumor types. For nine tumor types (DLBC, SKCM, ESSC, BRCA, COAD, KIRC, LIHC, LUSC and OV), lineage-specific dependency data were available. Factors that are lineage-specific dependencies in the relevant tumor type are indicated by bold font and an asterisk. Data curated from DepMap.org. (U) Percentage of tumor types with dependency (CERES) scores ≤ −0.4, −0.6, −0.8 or −1.0, on at least one candidate MTF in at least 50% of the corresponding cell lines.

### Candidate MTFs are targets of somatic mutations in cancer

We tested whether candidate MTFs were more likely to be somatically mutated in cancer than other TFs. We calculated the mutation rate across all coding exons and identified all significant genes (p < 0.05), using data for somatic single nucleotide variants from the PanCancer Analysis of Whole Genomes (PCAWG) project (https://icgc.org). Data were available for 21 (out of 34) cancer types represented in Figure 1E. We compared mutation burden across candidate MTFs to mutation burden of TFs with comparably high levels of expression (within the top 5% of all TFs; expression rank ≤ 79) but low CaCTS scores (CaCTS rank > 79). Candidate MTFs were more likely to be mutated than non-candidate MTFs that were also highly expressed in the same tumor type (p = 1.0 × 10^−4^, two-tailed Pearson’s Chi-squared test), particularly in BLCA, BRCA, KICH, LUSC and PAAD. (Table 1, Figure 3). Significantly mutated MTFs included factors previously known to be somatically mutated in a tumor-specific manner - *FOXA1* in BRCA and PRAD (Annala et al., 2018; Robinson et al., 2013) and *NFE2L2* in squamous lung tumors (Frank et al., 2018; Xiong et al., 2018). *TRPS1*, a known essential gene and lineage-specific factor in breast cancer was mutated in 11 out of 195 breast cancer cases (p = 4.5 ×1 0^-3^); other novel factors with significant mutation burdens included *TBX3* in bladder tumors (mutated in 5 out of 23 cases, p = 1.8 × 10^−3^) and *PAX8* in UCEC (mutated in 4 out of 44 cases, p = 0.01).

**Table 1.** Somatic mutations of candidate MTFs. MTFs with a significant burden of single nucleotide variants (SNVs) across coding exons, compared to all coding exons in the genome (listed in order of significance).

| Tumor Type | TF | Sample Size (n) | Mutated Samples (n) | SNVs (n) | P-value | Adjusted P-value |
| --- | --- | --- | --- | --- | --- | --- |
| LUSC | *NFE2L2* | 47 | 22 | 25 | $3.33 \times 10^{-16}$ | $2.00 \times 10^{-15}$ |
| PRAD | *FOXA1* | 275 | 11 | 11 | $5.13 \times 10^{-9}$ | $9.74 \times 10^{-8}$ |
| KIRC | *MAF* | 143 | 8 | 8 | $8.25 \times 10^{-4}$ | 0.01 |
| KICH | *FOXI1* | 43 | 2 | 2 | $1.46 \times 10^{-3}$ | 0.02 |
| BLCA | *TBX3* | 23 | 5 | 5 | $1.82 \times 10^{-3}$ | 0.03 |
| BRCA | *TRPS1* | 195 | 11 | 13 | $4.48 \times 10^{-3}$ | 0.04 |
| BRCA | *FOXA1* | 195 | 6 | 6 | $5.08 \times 10^{-3}$ | 0.04 |
| READ | *SOX9* | 52 | 9 | 11 | $6.15 \times 10^{-3}$ | 0.11 |
| COAD | *SOX9* | 52 | 9 | 11 | $6.15 \times 10^{-3}$ | 0.14 |
| UCEC | *MSX1* | 44 | 3 | 3 | $8.61 \times 10^{-3}$ | 0.09 |
| PAAD | *CREB3L1* | 234 | 5 | 5 | 0.01 | 0.07 |
| PAAD | *BHLHE40* | 234 | 5 | 5 | 0.01 | 0.07 |
| KICH | *MECOM* | 43 | 2 | 2 | 0.01 | 0.11 |
| UCEC | *PAX8* | 44 | 4 | 5 | 0.01 | 0.09 |
| BLCA | *ELF3* | 23 | 4 | 5 | 0.01 | 0.09 |
| BLCA | *ID1* | 23 | 2 | 2 | 0.01 | 0.09 |
| READ | *MYC* | 52 | 7 | 10 | 0.01 | 0.19 |
| COAD | *MYC* | 52 | 7 | 10 | 0.01 | 0.23 |
| ESAD | *AHR* | 97 | 7 | 7 | 0.01 | 0.32 |
| BRCA | *XBP1* | 195 | 4 | 7 | 0.01 | 0.18 |
| LIHC | *XBP1* | 324 | 6 | 6 | 0.01 | 0.47 |
| LIHC | *RORC* | 324 | 7 | 8 | 0.01 | 0.47 |
| BRCA | *GATA3* | 195 | 4 | 4 | 0.01 | 0.20 |

**Figure 3.**
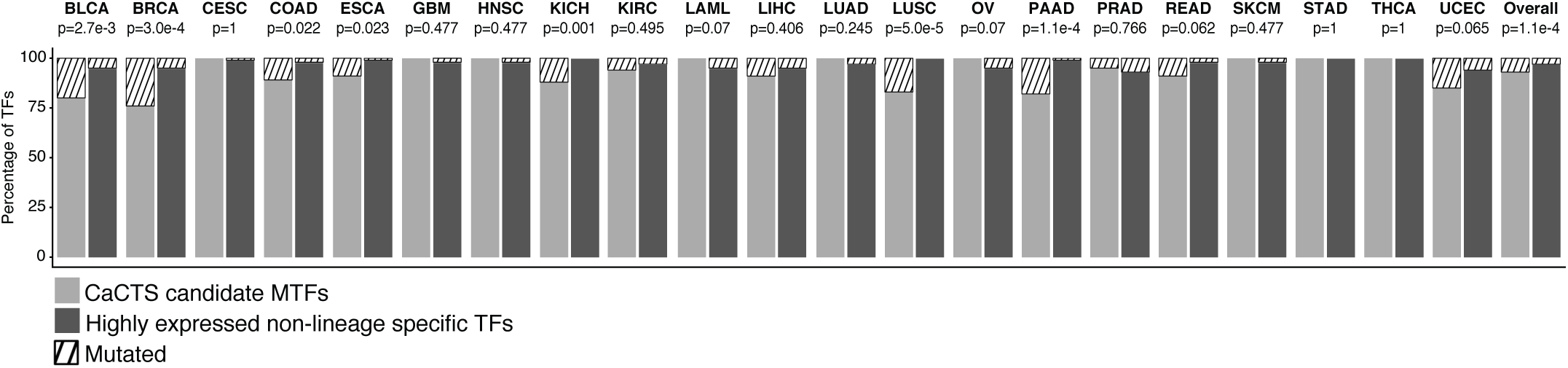
Candidate MTFs are frequently somatically mutated in relevant tumors. Proportions of candidate MTFs mutated in 21 tumor types. Highly expressed TFs (within the top 5% of all TFs) with low CaCTS scores were used as a comparison group. P-values indicate results from two-tailed Chi-squared tests.

### Subtype-specific MTF predictions reveal subtype-specific regulators

Many tumors characterized by a shared anatomic origin can be stratified into molecular and/or histologic subtypes. It has become increasingly clear that different subtypes often have markedly different prognoses and responses to therapy. Subtypes can also reflect different cells of origin and so we sought to identify candidate MTFs in tumors stratified into clinically relevant subtypes. Subtype annotation was available for 7,259 of the 9,691 tumors in our original data set; CaCTS scores were consistent when we reran CaCTS (for the 34 major tumor groups) using the smaller data set of 7,259 samples, indicating that the reduced set of samples is representative of the original data set (Figure S4A). We stratified our collection of tumors into subtypes based mostly on molecular features (expression, methylation, coding mutations or copy number alterations) except in the following three instances where we used histologic classifications: kidney chromophobe tumors (KICH) were divided into classic and eosinophilic histologic groups; the heterogeneous group of sarcomas (SARC), were divided into dedifferentiated liposarcomas, uterine leiomyosarcomas, soft tissue leiomyosarcomas, myxofibrosarcomas, undifferentiated pleomorphic sarcomas, malignant peripheral nerve sheath tumors and synovial sarcomas. Finally, uterine carcinosarcomas (UCS) were stratified into endometrioid-like and serous-like tumors, which was primarily based on somatic mutations but also corresponded to histologic classifications (Cherniack et al., 2017). To select the most clinically relevant subtype classifications for the remaining tumors we used TCGAbiolinks (Colaprico et al., 2016) (ACC, BCLA, BRCA, COAD, ESAD, GBM, HNSC, LGG, LIHC, LUAD, LUSC, OV, PRAD, READ, SKCM, STAD, THCA, UCEC) and primary publications from TCGA (CESC (Cancer Genome Atlas Research Network et al., 2017b), CHOL (Farshidfar et al., 2017), KICH (Davis et al., 2014), KIRC (Cancer Genome Atlas Research Network, 2013), KIRP (Cancer Genome Atlas Research Network et al., 2016), LAML (Cancer Genome Atlas Research Network et al., 2013), MESO (Hmeljak et al., 2018), PAAD (Cancer Genome Atlas Research Network, 2017), PCPG (Fishbein et al., 2017), SARC (Cancer Genome Atlas Research Network, 2017), TGCT (Shen et al., 2018), THYM (Lee et al., 2017), UCS (Cherniack et al., 2017), UVM (Robertson et al., 2017)). Subtype annotations were not available for DLBC and ESCC. Table S7 details how the 34 tumor groups were stratified into a total of 140 molecular and histologic subtypes. To implement the CaCTS algorithm, we queried the average expression of 1,578 TFs in each subtype (Table S8) against a background data set which contained the average TF expression in the other tumor subtypes, excluding all other tumors of the same major type (see Methods). The CaCTS algorithm identified a total of 439 different candidates across the 140 tumor subtypes; this included all candidates identified in our initial analyses plus 166 (38%) new factors that were only identified in the subtype-stratified analyses (Table S9). Subtype stratification had a major impact on the candidates identified for a number of tumor subtypes, particularly breast adenocarcinoma (BRCA), bladder urothelial carcinoma (BLCA) and cervical squamous cell carcinoma and endocervical adenocarcinoma (CESC) (Figure 4A). We compared candidate MTFs for the two predominant subtypes of breast cancer - luminal A (lumA, 55 % of BRCA cases) and basal (30% of cases). Candidate MTFs were vastly different between the two subtypes, with only two shared factors (*TRPS1* and *LTF*), consistent with existing evidence suggesting these two tumor types represent different cell states and cells-of-origin (Polyak, 2007). When we stratified DepMap dependency data for breast cancer cell lines based on the BRCA subtype of the cell line models, hierarchical clustering based on dependency across a union set of luminal A and basal BRCA candidates divided cell lines largely by subgroup (Figure 4B). Luminal A-specific candidates were not dependencies in basal-type cell lines, and *vice versa*. We detected some known subtype-specific factors including *GATA3* in the luminal A subtype (Ciriello et al., 2015; Shaoxian et al., 2017), and *FOXC1* and *SOX9* in triple negative breast cancer (which is enriched in the basal subtype) (Wang et al., 2015). Luminal A candidates largely overlapped with the candidates identified in the initial CaCTS analyses, which is to be expected as this comprised over 50% of the BRCA cohort. We also identified four additional candidate MTFs for luminal A breast cancer - *AEBP1, MYB, SREBF1* and *TBX3. TBX3* is recurrently mutated in luminal A tumors (Ciriello et al., 2015) but has not been implicated as an MTF, and the other factors also represent novel candidate MTFs for this tumor type. Only 2 out of 11 candidate MTFs identified for basal breast tumors were also candidates in the initial BRCA analyses (*LTF* and *TRPS1*). Novel candidates for basal BRCA included *NFIB (Moon et al., 2011)*, which has been previously implicated in epigenetic reprogramming during small cell lung cancer metastasis (Denny et al., 2016) and *CREB3L2*, which is commonly fused to *FUS* in low-grade fibromyxoid sarcomas (Matsuyama et al., 2006), but has not been studied in the context of basal-type breast cancer.

**Figure 4.**
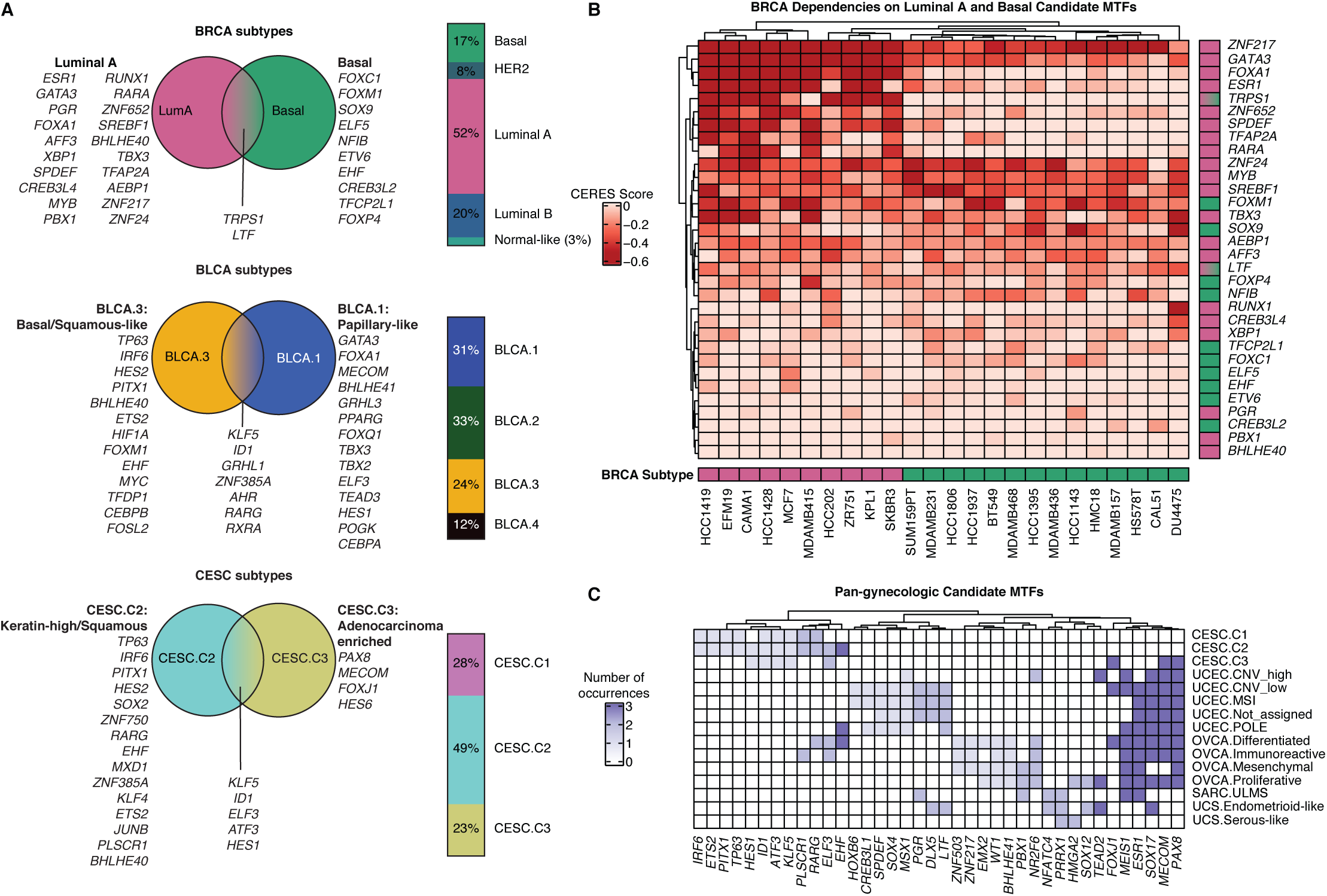
Diversity of candidate MTFs define tumor subtypes. (A) Distinct sets of candidate MTFs were identified for tumor subtypes in BRCA, BLCA and CESC. Stacked bar plots show frequency of molecular subgroups in each of the 3 tumor types. (B) Unsupervised clustering of luminal and basal breast cancer cell lines based on dependencies on a union set of Luminal A and Basal candidate MTFs. Cell lines cluster by molecular subtype. Green boxes correspond to Luminal A cell lines and Luminal A candidate MTFs; fuchsia denotes Basal/TNBC cell lines and candidate MTFs. (C) Candidate MTFs across pan-gynecologic tumor subtypes.

When we stratified molecular groups in bladder and cervical cancer we found that squamous-enriched group BLCA.3 and CESC “keratin” groups CESC.C1 and CESC.C2 have squamous differentiation TFs as candidates (*TP63, IRF6* and *PITX1*) and now cluster together with other squamous tumor types (LUSC, ESSC and HNSC). In contrast, adenocarcinoma subgroups from the same organs cluster more distantly (for example CESC.C3 clusters with uterine and ovarian adenocarcinoma) (Figure S4B) and have distinct MTF candidates. For BLCA.1 “papillary like” tumors, candidates include *GATA3* (Figure 4A), which is shared with BLCA.2 and BLCA.4, all breast cancer subtypes except basal, and the cortical admixture and pseudohypoxia subtypes of PCPG. *FOXA1* is a candidate MTF in BLCA.1 (but not in other bladder cancer subtypes), LUAD.2, all subtypes of prostate and non-basal breast cancer subtypes. *PAX8* and *MECOM* are candidates in CESC.C3, as well as OV and UCEC (Figure 4C). *FOXJ1* is also a candidate in CESC.C3, a factor that is crucial for ciliogenesis (Brody et al., 2000) and is also detected in the differentiated subgroup of OV, and the ‘copy number low’ subgroup of UCEC. Therefore, while squamous subgroups of bladder and cervix tumors cluster with squamous types from distant organs, adenocarcinomas share greater similarities with tumors derived from a similar developmental lineage.

Since *PAX8* and *MECOM* were candidates in CESC.C3 and OV, we hypothesized there may be additional factors shared across gynecologic tumor subtypes. We therefore compared MTF candidates in subtypes of cervical, uterine corpus endometrial and ovarian carcinomas, and uterine carcinosarcomas and leiomyosarcomas (now subdivided from the sarcoma metagroup). Overall six factors were common across two or more gynecologic tumor types - *PAX8, MECOM, SOX17, ESR1, MEIS1* and *FOXJ1. PAX8, MECOM* and *SOX17* were the top three shared candidate MTFs among this set of tumors (Figure 4C). Molecular subgroups of ovarian and uterine carcinoma (OV and UCEC) exhibited the greatest similarities, with CESC.C3, uterine leiomyosarcoma and uterine endometrioid-like sarcomas sharing 3, 2 and 2 candidate MTFs in common with the uterine/ovarian carcinoma metagroup, respectively. We also noted that for molecular subtypes of OV, MTF candidates largely mirrored those identified for OV as a whole, with a few modifications - the mesenchymal molecular subgroup of ovarian tumors uniquely lacked *SOX17* and *MECOM*, present in the other OV subtypes; *FOXJ1, BCL6, EHF* and *RARG* were candidates for differentiated tumors, but not the other subtypes; ELF3 was a candidate MTF for the differentiated and immunoreactive OV subtypes only; and proliferative-type tumors had four unique candidates - *HMGA2, SOX12, TEAD2* and *PLAGL2. HMGA2* is a known subtype-specific marker for the proliferative subgroup of OV (Cancer Genome Atlas Research Network, 2011) and TEAD family proteins likely cooperate with PAX8 to regulate gene expression in models of ovarian cancer (Adler et al., 2017; Elias et al., 2016). *SOX12* and *PLAGL2* are novel candidates for this subtype of OV tumors.

### Using candidate MTFs to build core regulatory circuitry models in ovarian cancers

To validate the CaCTS algorithm, we tested our success at identifying MTFs for OV. The TCGA OV study consists exclusively of high-grade serous ovarian cancers (HGSOCs). HGSOCs are relatively rare tumors for which MTFs are currently unknown, and novel therapeutic targets are urgently needed due to the frequent late-stage diagnoses and high rates of tumor recurrence. We identified 14 candidate MTFs for OV (listed in order of ranked CaCTS score): *WT1, EMX2, SOX17, MEIS1, BHLHE41, PAX8, ESR1, ZNF503, MECOM, TGIF2, NR2F6, PBX1, ZNF217* and *PLSCR1* (Figure 2S). PAX8 has not previously been characterized as an MTF, but is a known lineage-specific dependency in this tumor type (Cheung et al., 2011); and both PAX8 and WT1 are used as clinical biomarkers for serous ovarian carcinomas. We tested whether these factors show orthogonal characteristics of MTFs using ChIP-Seq experiments and *in vitro* knockdown studies.

Ten of these factors were associated with SEs in at least 4 (out of 12) tumor samples (Figure 5A,B; Figure S5) and eight factors - *MEIS1, SOX17, PAX8, WT1, ZNF217, BHLHE41, MECOM* and *PBX1* - were all associated with SEs in at least 11 of the 12 tumors. We decided to focus on *PAX8, SOX17* and *MECOM* as these are also pan-gynecologic factors and are marked by especially large SEs in primary ovarian tumors. We performed TF ChIP-seq in a HGSOC cell line (Kuramochi) and found these factors bind regulatory elements associated with the genes encoding these factors, consistent with their formation of a core regulatory circuit (CRC) (Figure 5C,D). Additionally, they bind at three clinical biomarkers of the disease: *WT1, MUC16* (CA125), and *HE4* (Köbel et al., 2008) (Figure 5E). CRC factors are also expected to drive expression programs by co-occupying enhancers across the genome, and we observed a consistent co-localization of these factors genome-wide (Figure 5F). Indeed the 2^nd^ and 13^th^ top ranked PAX8 binding peaks were at the *SOX17* gene locus (Figure 5G). Furthermore, proximity ligation assays confirmed that these proteins are part of the same complex (Figure 5H,I).

**Figure 5.**
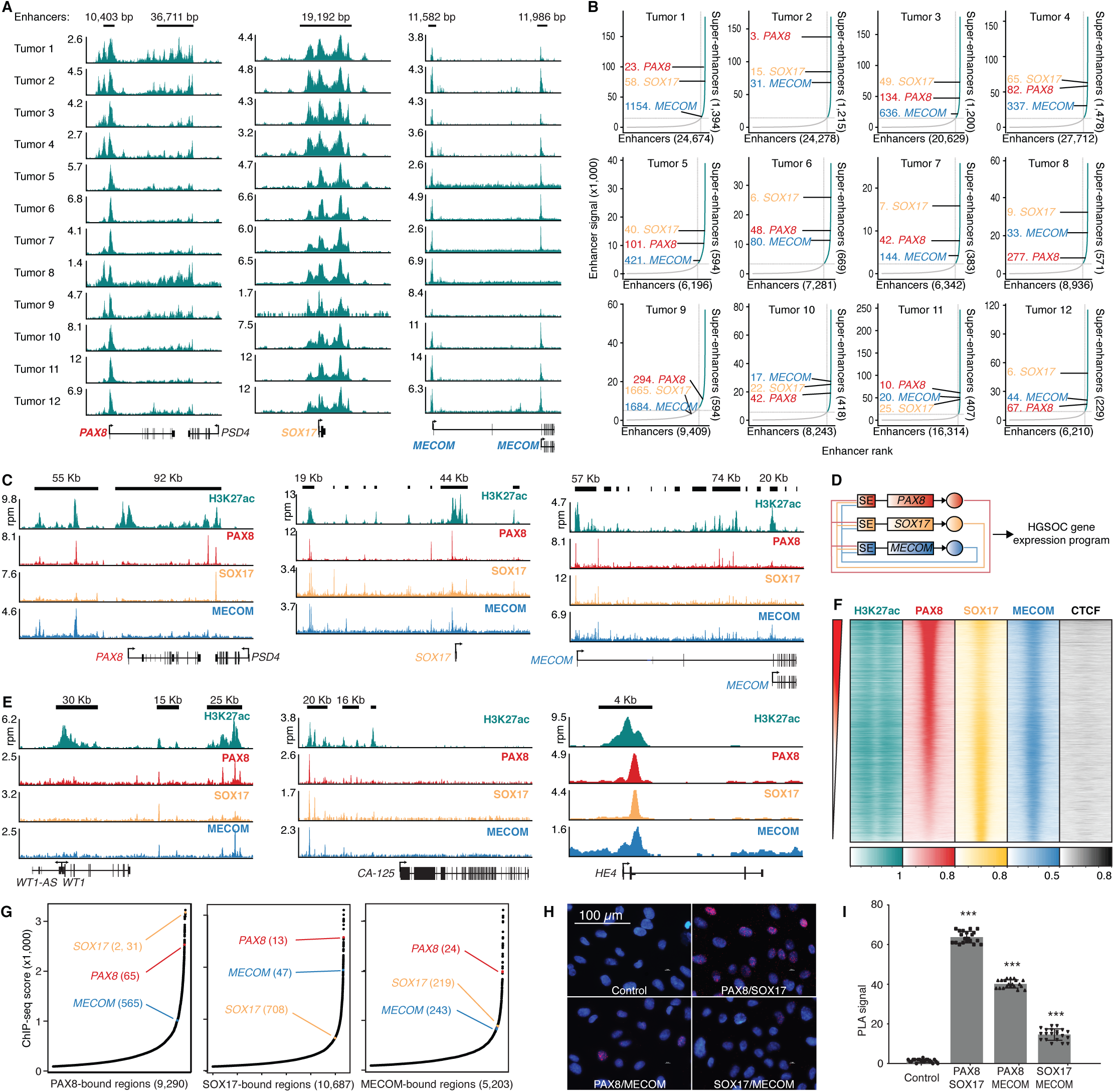
PAX8, SOX17 and MECOM are candidate MTFs in high-grade serous ovarian cancer (HGSOC). (A) PAX8, SOX17 and MECOM coincide with super-enhancers in primary ovarian tumors. H3K27ac ChIP-seq data were generated in 12 primary tumors, and data were normalized to counts per million (CPM) mapped reads. (B) SEs associated with PAX8, SOX17 and MECOM tend to be highly ranked. Stitched enhancers were ranked by H3K27ac ChIP-seq signal. (C) PAX8, SOX17 and MECOM occupy their own and each other’s SEs. Enhancers are indicated as black bars. (D) Proposed core regulatory circuit for HGSOC. (E) PAX8, SOX17 and MECOM bind to enhancers of HGSOC clinical biomarkers. (F) Global co-binding of PAX8, SOX17 and MECOM. Each row is a PAX8 ChIP-seq peak. CPM normalized ChIP-seq reads were plotted for a 4 kb window centered on each binding site. Rows are ordered by decreasing PAX8 signal. (G) PAX8, SOX17 and MECOM ChIP-seq peaks were ranked by CPM-normalized signal. Strong binding sites for each factor are detected proximal to their own and each other’s gene loci. (H) Proximity ligation assay performed in Kuramochi cells. Each red dot represents a single interaction. Nuclei were counter-stained blue with DAPI (blue). Scale bar: 100 μm. Image magnification × 40. (I) Quantified PLA signal per cell. In C, E, F and G, ChIP-seq data were generated in Kuramochi cells and normalized to counts per million (CPM) mapped reads.

To study the collaboration between PAX8, SOX17, and MECOM in driving tumor cell survival, we analyzed tumor cell survival in their absence. Overall 5 (out of 14, 36%) OV MTF candidates showed moderate to high levels of essentiality in at least one HGSOC cell line (minimum CERES score of −0.4 or less) (Figure 2S, 6A), particularly for PAX8 and MECOM where minimum CERES scores were −1.3 and −0.7, respectively. These two factors are also selective dependencies for OV (Cheung et al., 2011). PAX8, SOX17 and MECOM dependency correlates with level of expression, especially in ovary samples; the higher these factors are expressed the more likely they are to be essential. This is consistent with a model in which these factors are playing oncogenic roles (Figure 6A). In addition, *PAX8, SOX17* and *MECOM* gene loci are amplified in 6%, 11%, and 36% HGSOCs, respectively (Figure 6B). Using RNAi we depleted PAX8, SOX17, and MECOM protein expression by 80-95% and RNA expression by 80-95% (Figure 6C,D, Figure S6A). Following PAX8, SOX17, and MECOM knockdown, colony formation of HGSOC cells was reduced by 76% (standard deviation (s.d.) = 3.1%; p < 0.001), 44% (s.d. = 30%; p = 0.013) and 51% (s.d. = 17.9%; p = 0.004) respectively in comparison to siNT1 (Tukey’s Multiple Comparison Test, n=4) (Figure 6E). Comparisons to siNT2 were similar to that of siNT1 (Figure 6E). Together these data suggest a requirement for maintained PAX8, SOX17, and MECOM expression for HGSOC cell viability.

**Figure 6.**
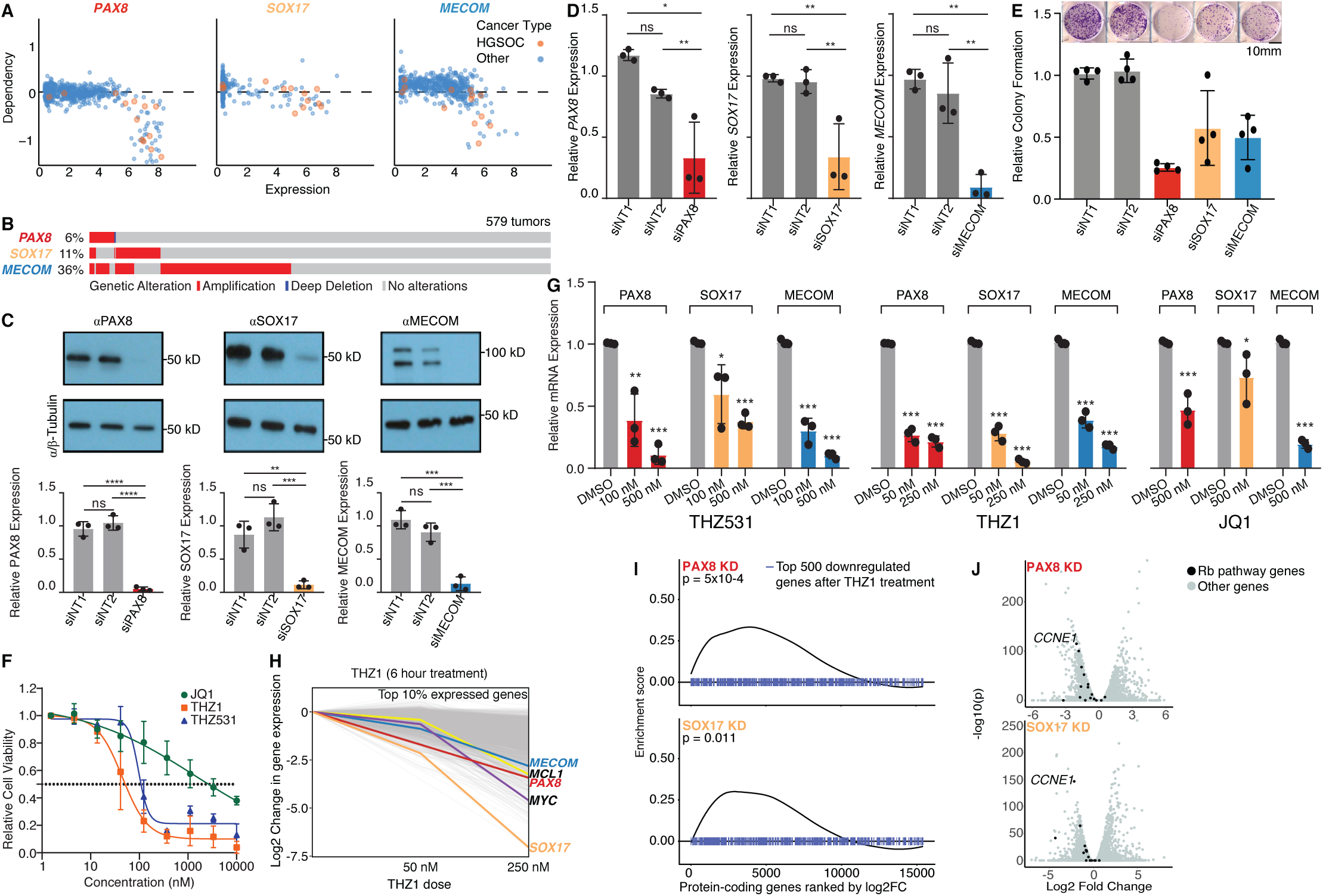
PAX8, SOX17 and MECOM are functional dependencies in high-grade serous ovarian cancer (HGSOC). (A) PAX8, SOX17 and MECOM are selective dependencies in HGSOC cell lines. (B) PAX8, SOX17 and MECOM are commonly amplified in HGSOC tumors. (C-E) Successful siRNA-mediated knockdown of PAX8, SOX17 and MECOM in OVCAR4 HGSOC cells. Knockdown was confirmed using (C) Western blot, plus (D) qRT-PCR. (E) Knockdown of PAX8, SOX17 and MECOM results in significantly reduced colony formation in anchorage independent growth assays. * p < 0.05, ** p < 0.01, *** p < 0.001; Students paired T-test; ns, not significant. Error bars indicate standard deviation of mean values from three independent experiments (performed with technical triplicates). (F) Dose response curves for OVCAR4 cells treated with a range of doses of THZ1, THZ531 and JQ1 for 72 hours. Control cells received vehicle (DMSO). Non-linear fit curves shown. Data are representative of three independent experiments. (G) Quantification of PAX8, SOX17 and MECOM expression by qRT-PCR, following a 6 hour treatment with THZ1, THZ531 and JQ1. Gene expression for each factor was normalized to the average expression *ACTB* and *GAPDH*, and expression fold change for drug treatments calculated relative to vehicle-treated control cells. * p < 0.05, ** p < 0.01, *** p < 0.001; Students paired T-test; ns, not significant. Error bars indicate standard deviation of mean values from three independent experiments (performed with technical triplicates). (H) PAX8 and SOX17 are among the most sensitive genes in THZ1-treated Kuramochi cells. (I) Gene set enrichment analysis of the top 500 downregulated genes following PAX8 and SOX17 knockdown, compared to a ranked list of THZ1 responsive genes. (J) Global analyses of PAX8 and SOX17 target genes, identified by performing RNA-seq 72h after knockdown. Rb pathway genes are among the most significantly downregulated transcripts.

### Targeting an MTF-driven oncogenic expression program in HGSOC

General transcription inhibitors are showing remarkable anti-cancer effects across multiple cancer types, which is thought to be largely due to their preferential activity towards MTFs (Delmore et al., 2011; Durbin et al., 2018; Wang et al., 2015). High-grade serous ovarian cancer models are exquisitely sensitive to pharmacologic inhibition of CDK7/12/13 (Francavilla et al., 2017; Kwiatkowski et al., 2014; Zeng et al., 2018) and many cell lines are also sensitive to inhibition of BET family bromodomain proteins with JQ1 (Baratta et al., 2015). Consistent with these results, OVCAR4 HGSOC cells exhibit notably low IC50 values of 45 nM for THZ1 and 1.3 μM for JQ1. OVCAR4 cells are also sensitive to THZ531, a covalent inhibitor of cyclin dependent kinases 12 and 13, again with IC50 values in the nanomolar range (97 nM) (Zhang et al., 2016) (Figure 6F). The three candidate MTFs were exquisitely sensitive to these molecules (Figure 6G, Figure S6B). PAX8 and SOX17 were particularly sensitive to THZ1 treatment and are among the 10%-most sensitive among highly expressed protein-coding transcripts in low-dose (50 nM) treatment with THZ1 (Figure 6H). The potent inhibition of these factors with low-dose treatment is most relevant in terms of target engagement and the concentration range that selectivity is observed (Kwiatkowski et al., 2014). Finally, RNA-seq revealed that PAX8 and SOX17 knockdowns both phenocopy effects of low-dose THZ1 treatment, suggesting these factors, at least in part, explain the anti-cancer effect of this drug in ovarian cancer cells (Figure 6I). PAX8 and SOX17 target genes are largely overlapping, with some of the most downregulated genes falling into cell cycle, DNA replication, and DNA division pathways (Figure S6C,D), including cell-cycle regulators in the retinoblastoma pathway such as known ovarian cancer oncogene *CCNE1* (Cancer Genome Atlas Research Network, 2011; Karst et al., 2014; Kuhn et al., 2016; Patch et al., 2015) (Figure 6J).

## Discussion

Core regulatory circuitries (CRCs) represent a potentially universal vulnerability in tumor cells, and consequently, they are likely to represent potent therapeutic opportunities for many cancer types. Given recent developments in targeting CRCs through the use of general transcription inhibitors, this approach to prioritizing candidate master regulators based solely on RNA-seq data is timely, particularly for tumor types where limited access to tumor specimens prohibits the generation of the ChIP-seq data typically required to identify candidate MTFs. The candidate MTFs recovered by the CaCTS algorithm include both known and novel MTFs, which are especially valuable for tumor types in which transcriptional circuits are poorly characterized. As a proof-of-concept we performed functional validation and confirmed MTF features for PAX8, SOX17 and MECOM in HGSOC cells; demonstrating that the CaCTS predictions recovers novel critical regulators. In the DepMap cell line dependency data we observed striking clustering patterns within each tumor type, with factors and cell lines clustering by co-dependencies for many tumor types, demonstrating the success of the CaCTS approach to identify transcriptional circuitries. Cell lines that show the greatest dependence on tumor-specific MTFs may be the superior models to use for translational studies; for example in HGSOC, Kuramochi, OAW28 and ONCODG1 exhibited similar dependencies on candidate MTFs, and have all been prioritized as cell lines models that faithfully recapitulate molecular hallmarks of HGSOC (Domcke et al., 2013).

We predicted candidate factors for major tumor groups, as well as clinically relevant molecular and histologic subtypes. This analysis revealed some common factors across diverse origins, a phenomenon most clearly illustrated by squamous tumors where tumors across five diverse anatomic sites shared three common factors (*TP63, KLF5* and *IRF6*). A further three factors were shared by 3 sites - *RARG* (BLCA, CESC and HNSC), *PITX1* (CESC, ESSC and HNSC) and *NFE2L2* (ESSC, HNSC and LUSC). *KLF5* is shared between squamous tumors and adenocarcinomas of the esophagus, cervix and bladder suggesting this factor may have different targets in the different tumor types. In some instances, squamous tumors have both squamous and organ-specific candidate MTFs, such as *ID1* and *KLF5* in cervix and *AHR* and *KLF5* in esophageal tumors; suggestive of dual circuitries with cooperating factors dictating lineage-specific and functional programs. Using CaCTS we were able to make MTF predictions for rare and common tumor types that lack publicaly available SE or functional dependency data. We were unable to retrieve high-quality H3K27ac data to call SEs for 14 out of the 34 major tumor types (ACC, CHOL, HNSC, KICH, KIRP, LUSC, MESO, STAD, TGCT, PCPG, THCA, THYM, UCS, UVM) and for five additional tumor types (CESC, LUAD, SARC, ESAD and ESSC), primary tumor data were not available and so we used ChIP-seq data from cell line models, which may not closely recapitulate the epigenetic signatures of disease. In addition, 6 tumor types are not represented in DepMap (ACC, KICH, KIRP, TGCT, THYM, UCS) and there are limited dependency data (three cell lines or fewer) for four other tumor types (CESC, CHOL, THCA and UVM). Overall, around half of the tumor types represented in our study currently have limited publicly available data to predict MTFs, and for six tumor types neither H3K27ac *nor* dependency data are available - adrenocortical carcinomas, kidney chromophobe tumors, kidney renal papillary cell carcinomas, testicular germ cell tumors, thymoma and uterine carcinosarcomas, and so MTF prediction is not possible based on current methods. Discussion of the MTFs identified for these tumor types are included as a Supplementary Note.

Cancer MTFs represent attractive therapeutic targets, given their essential role in governing cell state, and the ‘transcriptional addiction’ phenomenon (Bradner et al., 2017), whereby cancer cells become highly dependent on maintained high level expression of a select handful of MTFs. Consistent with this, many of the candidate MTFs identified by CaCTS were lineage-specific essentialities in the relevant tumor type. General transcriptional inhibitors, which show preferential activity towards MTFs, offer an efficient approach to anti-cancer treatment, rather than developing drugs to target each individual MTF. A number of such agents are now in clinical trials. Functional validation of the OV candidate MTFs, PAX8, SOX17 and MECOM, revealed that each were sensitive to general transcription inhibition, with PAX8 and SOX17 particularly vulnerable to CDK7 inhibition with THZ1. This suggests that PAX8 and SOX17 contribute to anti-proliferative effects of this drug, in addition to its previously studied effects on MYC and MCL1 (Zeng et al., 2018). As PAX8, SOX17 and MECOM are shared factors in other tumor types and subtypes, these results may be applicable to other tumor types dependent on these factors, including gynecologic tumor subtypes that share the greatest similarities with OV (UCEC and CESC C3) and some non-gynecologic tumors - thyroid and kidney carcinomas (THCA, KIRP, KICH and KIRC) where *PAX8* is a candidate MTF, and esophageal adenocarcinomas, kidney chromophobe tumors and stomach adenocarcinomas where *MECOM* is a candidate. Early downregulated genes following PAX8 and SOX17 depletion included genes in the Rb pathway, indicating a mechanism of action for anti-cancer activity of THZ1 in ovarian cancer models. Critically, ovarian cancer oncogene *CCNE1* was downregulated −2.0 and −2.2 fold following PAX8 and SOX17 depletion, respectively, consistent with *CCNE1* being a target of these MTFs. Rb pathway alterations predict responses of patient derived xenograft models to SY-1365 (Hu et al., 2019), a covalent CDK7 inhibitor currently in clinical trials for advanced breast and ovarian cancer (NCT03134638). While we validated 3 factors in OV, additional MTFs may contribute to the transcriptional circuitry of HGSOC. One such factor is WT1, an ovarian cancer biomarker, whose SE was co-bound by PAX8, SOX17 and MECOM. *MEIS1* was another candidate, this factor was associated with SEs in all 12 tumors and was the highest-ranking SE-associated MTF for this tumor type. Cells dependent on PAX8 also tended to be dependent on ZNF217, a factor implicated in breast cancer (Frietze et al., 2014; Littlepage et al., 2012). Other shared factors with breast cancer included ESR1 (known to interact with ZNF217) and PBX1. Compared to PAX8 and SOX17, MECOM was less responsive to THZ1 and THZ531 (although MECOM expression was highly sensitive to JQ1), and as a candidate this factor may not fully fit the canonical MTF model. *MECOM* was the lowest ranking of the three factors based on SE associations, and was least lineage-specific of the three, as it is shared in many adenocarcinomas. We noted for OV and the majority of tumor types we examined, that the tumor MTF predictions were very similar to those previously predicted for normal tissues of the same organ (D’Alessio et al., 2015) suggesting that a major mechanism of tumorigenesis involves aberrant reinforcement of developmental transcriptional programs, or that normal MTF activities become perturbed to acquire oncogenic properties during cancer development. MTFs predicted by CaCTS are more likely to contain somatic mutations than TFs with comparably high levels of expression but lacking lineage-restricted patterns of expression. This observation supports the idea that somatic mutations that bestow pro-oncogenic properties on developmental MTFs are selected for during cancer development. We present a prioritized collection of candidate MTFs for 34 major tumor types and 140 tumor subtypes, which will be enriched for *bone fide* MTFs, but it will also likely contain false positive. For users of this resource, we recommend integration of complementary data, where available - such as SE landscapes, dependency data (Rauscher et al., 2019), and motif-based circuitry mapping (Federation et al., 2018) to inform the design of functional validation experiments. We note that not all cancer MTFs will fulfill all the canonical MTF criteria - for example *ZNF217* has lower levels of expression in basal-type breast tumors (expression rank = 232), but high levels of dependency (minimum dependency in basal BRCA cell lines = −1.12). An additional caveat to this approach is that nomination of candidates is directly related to the composition of the background data set. Overall TF expression across several of the tumor types were quite similar (for example, in the kidney tumor types). As such, it is possible that additional tumor type-specific factors exist but were lowly ranked. Despite this, known regulators were efficiently retrieved with this analysis, suggesting our background data set was sufficiently heterogenous to minimize this issue. In addition, our analyses were restricted to the pan-cancer data set compiled by TCGA, so some tumor types are not represented. Metastatic and treatment resistant tumor states, where new therapies are most urgently needed, are largely absent.

In closing, we present a timely and valuable resource of candidate MTFs for tumor types where transcriptional circuitries are currently unknown, and leverage this resource to identify PAX8, SOX17 and MECOM as master regulators for aggressive high-grade serous ovarian cancers.

## Acknowledgements

This project was supported by an Ovarian Cancer Research Fund Alliance Liz Tilberis Early Career Award (599175) (K.L.), Ovarian Cancer Research Fund Alliance Program Project Development (373356) (B.Y.K., K.L.), the American Lebanese Syrian Associated Charities (B.J.A.), a Southern California Clinical and Translational Science Institute Core Voucher (V148) (K.L., R.I.C.) and grants CA155258 and CA213333 (R.A.Y.). The research described was supported in part by NIH/National Center for Advancing Translational Science (NCATS) UCLA CTSI Grant Number UL1TR001881. R.N. is supported in part by a Ruth L. Kirschstein Institutional National Research Service Award (T32) from the NIH (grant number 5 T32 GM 118288-2). Please add the following grant support and disclosures. I.A.K. was supported by an American Society of Clinical Oncology Young Investigator Award, an American Cancer Society Postdoctoral Fellowship, and an Ovarian Cancer Research Alliance Mentored Investigator Award. IAK is a consultant and shareholder of Dewpoint Therapeutics.

This work used equipment and services provided at the Cedars-Sinai Biobank and Translational Research Core and the Applied Genomics, Computation and Translation (AGCT) Core. The results shown here are in whole or part based upon data generated by the TCGA Research Network: https://www.cancer.gov/tcga and the ICGC’s Pan-Cancer Analysis of Whole Genomes (PCAWG; https://pcawg.icgc.org/pcawg) consortium. Ovarian tumor specimens were collected as by the Jean Richardson Gynecologic Tissue and Fluid Repository at the University of Southern California, as part of the Women’s Cancer Program at Cedars-Sinai Medical Center, or as part of the Dana-Farber Cancer Institute ovarian tumor banking protocol.

## Disclosures

R.A.Y. is a founder and shareholder of Syros Pharmaceuticals, Camp4 Therapeutics, Omega Therapeutics, and Dewpoint Therapeutics. I.A.K. is a consultant and shareholder of Dewpoint Therapeutics.

## Online Methods

### Computational Methods

#### The Cancer Core Transcription factor Specificity (CaCTS) algorithm

PanCancer TCGA RNA sequence level 3 normalized data were downloaded from the GDC Data Portal using TCGAbiolinks functions GDCquery, GDCdownload and GDCprepare importing into R (http://www.r-project.org) for further analysis (Colaprico et al., 2016). Table S1 contains the tumor IDs for all the samples included in our analysis. After exclusion of recurrent, metastatic and non-tumor tissues, a total of 9,691 samples across 34 tumor types were available. Sample annotations were curated from TGCA publications (Cancer Genome Atlas Research Network. Electronic address: and Cancer Genome Atlas Research Network, 2017; Cancer Genome Atlas Research Network. Electronic address: and Cancer Genome Atlas Research Network, 2017; Cancer Genome Atlas Research Network, 2013; Cancer Genome Atlas Research Network et al., 2013, 2016, 2017b; Cherniack et al., 2017; Davis et al., 2014; Farshidfar et al., 2017; Fishbein et al., 2017; Hmeljak et al., 2018; Lee et al., 2017; Robertson et al., 2017; Shen et al., 2018)and TCGAbiolinks (Colaprico et al., 2016; Mounir et al., 2019) http://bioinformaticsfmrp.github.io/TCGAbiolinks/subtypes.html. Tumor types/subtypes included were ACC - Adrenocortical carcinoma, BLCA - Bladder Urothelial Carcinoma, BRCA - Breast invasive carcinoma, CESC - Cervical squamous cell carcinoma and endocervical adenocarcinoma, CHOL - Cholangiocarcinoma, COAD - Colon adenocarcinoma, DLBC - Lymphoid neoplasm diffuse large B-cell lymphoma, ESCA - Esophageal carcinoma, GBM - Glioblastoma multiforme, HNSC - Head and neck squamous cell carcinoma, KICH - Kidney chromophobe, KIRC - Kidney renal clear cell carcinoma, KIRP - Kidney renal papillary cell carcinoma, LAML - Acute myeloid leukemia, LGG - Brain lower grade glioma, LIHC - Liver hepatocellular carcinoma, LUAD - Lung adenocarcinoma, LUSC - Lung squamous cell carcinoma, MESO - Mesothelioma, OV - Ovarian serous cystadenocarcinoma, PAAD - Pancreatic adenocarcinoma, PCPG - Pheochromocytoma and Paraganglioma, PRAD - Prostate adenocarcinoma, READ - Rectum adenocarcinoma, SARC - Sarcoma, SKCM - Skin cutaneous melanoma, STAD - Stomach adenocarcinoma, TGCT - Testicular germ cell tumors, THCA - Thyroid carcinoma, THYM - Thymoma, UCEC - Uterine corpus endometrial carcinoma, UCS - Uterine carcinosarcoma, UVM - Uveal Melanoma. For the main analyses we preserved all grouping defined by TCGA, apart from for ESCA, which we divided into ESAD - Esophageal adenocarcinoma and ESSC - Esophageal squamous cell carcinoma.

Published lists of transcription factors (TF) were retrieved from Saint-Andre *et al*., (1,253 TFs) (Saint-André et al., 2016) and Lambert *et al*., (1,639 TFs) (Lambert et al., 2018). Merging both lists created a catalogue of 1,671 unique TFs, of which 1,578 were expressed in the pancancer data set. To calculate a Jensen-Shannon Divergence (JSD) score for each TF for each tumor type, we shifted the normalized expression values such that the new minimum normalized expression value is equal to 0. We quantified the specificity between the TF expression across different tumor types by creating two same-sized vectors to represent the observed and ideal pattern of lineage-specific TF expression. For the ‘observed pattern’, the vector was formed by values from the expression mean profiles of the query tumor type and the background data set. For instances where TF mean expression is equal to zero, we substituted 0 with 0.1^-17^ (as zero values are not compatible with the JSD calculations). Each element in this vector was divided by the sum of all elements such that the sum of the vector is now equal to one. For the ‘idealized pattern’, the vector was formed by a value of 1 at the position equivalent to that of the query tumor type and zeroes at all other positions. Using these two vectors, the JSD function was performed using the R package jsd version 0.1 and a specificity score was obtained for each TF, for each tumor type. The CaCTS score is the negative log10 of the JSD score. The final candidate MTF list for a given cancer type was defined by considering the intersection of the 5% most highly expressed TFs (expression rank ≤ 79) in said tumor type and the TFs in the top 5% when ranked by the CaCTS score.

#### Identification of super-enhancer associated genes

We collected publicly available H3K27ac ChIP-seq data from the Gene Expression Omnibus (https://www.ncbi.nlm.nih.gov/geo/) using search term “H3K27ac” “TCGA study abbreviation” or “tumor type” e.g. “BRCA” or “breast cancer”. We prioritized data generated for primary tumor tissues and only included data for cell lines when primary tumor data were not available. For OV, ESCA, PRAD, KIRC, GBM we used in-house tumor tissue H3K27ac ChIP-seq data. Data were processed using ENCODE pipeline version v1.2.0 and v1.1.7. ENCODE performed the alignment using bwa and peak calling MACS2. We only included samples that passed the following QC thresholds: total IP reads > 15 million; NSC > 1.05; RSC > 0.8; and FRiP > 0.1. The full curated list of H3K27ac ChIP-seq data sets used in this study can be found in Table S4. SE (SE) calls were obtained using the Rank Ordering of SE (ROSE2) algorithm for all tumor types, except prostate and kidney, where ROSE was used (Whyte et al., 2013). For ROSE2, we aligned to genome build hg19, with the following parameters: stitching distance -s 12500 and distance from TSS to exclude -t 2500. We selected SEs assigned to known transcription factors (Lambert et al., 2018; Saint-André et al., 2016).

#### Assessing enrichment of super-enhancer associated genes in CaCTS ranked lists

We implemented the gene set enrichment (GSEA) analysis using the R package FGSEA version 1.10, to evaluate the enrichment of SE associated genes with the list of TFs ranked by CaCTS scores for each tumor type. We applied the fgsea function with the parameter nperm equal to 10000. Numbers of SEs detected in each tumor are listed in Table S4. Some samples did not have SE-defining data sets available, so, given the biologic similarities of GBM and LGG, GBM SEs were used as a proxy to evaluate LGG MTFs predicted by CaCTS. Similarly, GSEA was performed on the READ candidates using COAD SEs.

#### Hierarchical clustering using CaCTS score

Clustering was performed using the *spearman* method and *complete* distance parameters. We selected a height cut-off equal to 0.63 to define the clusters, which resulted in 11 groups (in the 34 tumor group analyses). To compare our clusters with groups defined by TCGA (Hoadley et al., 2018) CaCTS clusters were matched to the TCGA cluster to which 50% (or more) of the samples were assigned.

#### Analyses of CaCTS TF dependencies

For the dependency analysis, we manually searched the Cancer Dependency Map Project database (Meyers et al., 2017) (DepMap Achilles 19Q1 public release - https://figshare.com/articles/DepMap_Achilles_19Q1_Public/7655150) for cell lines that rightfully correspond to the 34 tumor types. We found dependency data for 20 out of 34 (58.8%) cancer types, and retrieved 434 cell lines across these, with LUAD having the largest number of lines (n = 31) and CHOL having the lowest (n = 1). We performed hierarchical clustering of cell lines and the CaCTS TFs for the corresponding tumor type (method = ward.d2, distance = euclidean). We also calculated the percentage of tumor types with at least 1 predicted candidate with a CERES score < −0.4, < −0.6, < −0.8 or < −1 in at least 50% of the cell lines for that tumor type. Search terms used for each tumor type (for primary disease and subtype) were: ACC: “Adrenal Cancer”; BLCA: “Bladder Cancer”; LGG: “Brain Cancer”, then filtered by “Astrocytoma”,”Astrocytoma, anaplastic”, “Glioma, Neuroglioma”, “Oligodendroglioma”, “Oligodendroglioma, anaplastic”; BRCA: “Breast Cancer”; CESC: “Cervical Cancer”; CHOL: “Bile Duct Cancer”; COAD: “Colon/Colorectal Cancer”; ESCA-Adeno: “Esophageal Cancer” then filtered by “Adenocarcinoma”; ESCA-Squamous: “Esophageal Cancer” then filtered by “Squamous Cell Carcinoma”; GBM: “Brain Cancer” then filtered by “Glioblastoma”; HNSC: “Head and Neck Cancer”; KIRC: “Kidney Cancer” then filtered by “Renal Carcinoma, clear cell”, “Renal Adenocarcinoma, clear cell”; LIHC: “Liver Cancer”; LUAD: “Lung Cancer” then filtered by “Non-Small Cell Lung Cancer (NSCLCL), Adenocarcinoma”; LUSC: “Lung Cancer” then filtered by “Non-Small Cell Lung Cancer (NSCLCL), Squamous Cell Carcinoma”; DLBC: “Lymphoma” then filtered by “Diffuse Large B-cell Lymphoma (DLBCL)”; MESO: “Lung Cancer” then filtered by “Mesothelioma”; LAML: “Leukemia” then filtered by “AML”; PAAD: “Pancreatic Cancer”; PRAD: “Prostate Cancer”; COAD: “Colon/Colorectal Cancer”; SARC: “Sarcoma” then filtered by “Liposarcoma”; SKCM: “Skin Cancer” then filtered by “Melanoma”, “Melanoma, amelanotic”; STAD: “Gastric Cancer”; TGCT: “Embryonal Cancer”; THCA: “Thyroid Cancer”; UCS: “Endometrial/Uterine Cancer” then filtered by “Endometrial Stromal Sarcoma”; UCEC: “Endometrial/Uterine Cancer” then filtered by “Uterine/Endometrial Adenocarcinoma”, “Endometrial Carcinoma”, “Endometrial Adenocarcinoma”; UVM: “Eye Cancer”; PCPG: “Neuroblastoma”.

For OV and BRCA subtypes we manually curated cell lines for inclusion, as follows, OV: (OAW28, COV318, KURAMOCHI, SNU8, ONCODG, JHOS4, JHOS2, OVCAR8, COV504, COV362, OV90, X59M, OVCAR5, CAOV3, EFO21, A2780), BRCA (luminal A): (EFM19, HCC1428, CAMA1, HCC1419, MCF7, MDAMB415, SKBR3, ZR751, KPL1, HCC202, HMC18) BRCA (luminal B): (EFM19, HCC1428, CAMA1, HCC1419, MCF7, MDAMB415, SKBR3, ZR751, KPL1, HCC202, HMC18); BRCA (basal/TNBC): (MDAMB468, HCC1806, HCC1395, MDAMB436, MDAMB231, SUM159PT, BT549, HCC1937, MDAMB157, CAL51, HS578T, HCC1143, DU4475, HMC18); BRCA (HER2): (AU565, MDAMB453, JIMT1, HCC1954).

#### Tumor subtype MTF predictions

To predict candidate MTFs specific for each of the 140 tumor subtypes (Table S7), we implemented the same workflow developed for the 34 TCGA cancer types, however, instead of adding all samples for a query cancer type we selected one subtype at once to be the query group and all other cancer subtypes to be the background. For example, considering the four molecular subgroups for ovarian cancer we select ‘proliferative’ samples as query and all other 136 cancer subtypes as background, leaving out other molecular subgroups for OV, i.e. mesenchymal, differentiated and immunoreactive.

#### Somatic mutation analyses

We used coding single nucleotide variants (SNVs) from 2,715 tumors from the PCAWG project (https://icgc.org). We removed all SNVs that fall into regions of low mappability (wgEncodeDacMapabilityConsensusExcludable.bed). To identify frequently mutated genes, we calculated a background mutation rate for each sample. Let *X*_*i*_(*X*_*i*_ ∈ [0, *n*]) be a random variable that represents the number of samples with at least one mutation in the *i*th gene (where n is the total number of samples of a given tumor type), then *X*_*i*_ follows a Poisson binomial distribution (PBD) with a vector of probabilities 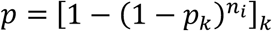, where *n*_*i*_ is the size of the coding sequence of the *i*th gene in base pairs, and *p*_*k*_ is the global background rate of sample *k* (*k* ∈ [1, *n*]) empirically estimated by the ratio of the total number of SNVs in sample *k* (*n*_*k*_) over the total coverage of all exons (in base pairs) (*n*_*cov*_): 

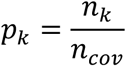

To determine whether the observed number of mutated samples in the *i*th gene, we calculated the probability of having at least *s*_*i*_ samples mutated, i.e., p-value_i_=*P*(*X*_*i*_ ≥ *s*_*i*_). P-values were adjusted used the Benjamini-Hochberg method.

#### Oncoplots of PAX8, SOX17, MECOM genetic aberrations

Data were obtained from cBioportal (Cerami et al., 2012; Gao et al., 2013). Data included 579 patients/samples with Ovarian Serous Cystadenocarcinoma, from the study TCGA Provisional.

### Experimental Methods

#### H3K27ac chromatin immunoprecipitation of primary HGSOC tissues

All tissues used were collected with informed consent and the approval of the institutional review boards of the University of Southern California, Cedars-Sinai Medical Center (CSMC), the Whitehead Institute for Biomedical Research (WIBR) and the Dana-Farber Cancer Institute (DFCI). All specimens profiled were primary, chemotherapy-naïve, high-grade serous ovarian cancers. Tumors 1-4 were profiled at CSMC and have been previously described (Corona et al., 2019). Briefly, 5 mm punches of OCT-embedded, pathology reviewed tumor specimens were taken from epithelial enriched regions. Tissues were subjected to ChIP-seq using an anti-H3K27ac antibody (DiAGenode, C15410196, Denville, NJ). Tumors 5-12 were profiled at WIBR. Thirty 30 μm sections of frozen tissues with a >90% tumor enrichment were washed with PBS and crosslinked with 1% formaldehyde for 10 minutes and quenched with 0.125 M glycine for 5 min at room temperature. Cross-linked material was resuspended in 1 ml lysis buffer (0.1%SDS, 1X TX-100, 10mM Tris-HCl pH=8, 1 mM EDTA pH=8, 0.1% Sodium Deoxycholate, 0.25% Sarkosyl, 0.3 mM NaCl, 1X PIC and 5mM Sodium butyrate) and sonicated for 20 minutes with a Covaris E220 instrument (10% duty cycle, 175 Peak Incident Power, 200 Cycles per burst, 1 mL AFA Fiber milliTUBEs). 8 μg soluble chromatin was immunoprecipitated with 10 μg of H3K27ac (Diagenode C15410196, lot# a1723-0041d) antibody. ChIP-seq libraries were constructed using Accel-NGS 2S DNA library kit from Swift Biosciences. Fragments of the desired size were enriched using AMPure XP beads (Beckman Coulter). 36-bp paired-end reads were sequenced on a Nextseq instrument (Illumina). Data were processed using the ENCODE pipeline, as described above, with SEs identified using ROSE.

#### Cell Culture

OVCAR4 and Kuramochi cells were cultured in RPMI-1640 supplemented with 10% FBS, 1× NEAA, 11.4 μg/ml of insulin, and 1× penicillin/streptomycin and maintained at 37°C with 5% CO_2_. Cells were passaged with 0.05% trypsin using standard cell culture procedures. Cells were confirmed to be negative for *Mycoplasma*, and were authenticated by profiling of short tandem repeats using the Promega Powerplex 16HS assay, performed at the University of Arizona Genomics Core (Table S10).

#### RNA interference and colony formation assays

OVCAR4 cells were reverse transfected with non-targeting (Dharmacon ON-TARGETplus Non-targeting Control Pool [NT1] and a second custom control pool containing: D-0012-03, D-001210-04, D-001210-05 [NT2]) or pooled PAX8, SOX17, and MECOM oligonucleotides (Dharmacon L-003778-00-0005, L-013028-01-0010 and L-006530-02-0005) by incubating 120 nM of each siRNA pool in Opti-MEM I (Thermo Fisher Scientific) for 5 minutes, which was then combined with a mix of Opti-MEM I and lipofectamine RNAiMAX (Thermo Fisher Scientific) and incubated for 20 minutes at room temperature. The transfection reagent mix was then combined with 300,000 cells and seeded in a 60mm dish. Media was replenished after 24 hours, and transfected cells used for analysis or assays 48 hours later. For colony formation assays, transfected cells were trypsinized, counted and 1,000 cells per condition were seeded in 6-well plates, in triplicate. Media were replenished once per week, and after 14 days, the cells were washed with 1× PBS (Thermo Fisher Scientific) three times and fixed with 10% formalin (Mckesson) for 20 minutes. Plates were then washed with water and stained with 0.1% crystal violet for 30 minutes. Excess crystal violet was washed with water and colonies counted manually.

#### Western Blotting

Cells were lysed with 100 μl of cell lysis buffer per 1 million cells (10mM HEPES, pH7.5 by KOH, 300mM NaCl, 0.1% NP-40, 5mM EGTA, with 10 μg/mL aprotinin, 10 μg/mL leupeptin, 1× protease inhibitor cocktail (Roche), 1× PhosSTOP Protease Inhibitor Cocktail (Roche), 1× PMSF (Sigma-Aldrich) and Supraise-in (Ambion) at 4°C for one hour. Lysed samples were then centrifuged at 4°C, 12,000 *g* for 10 minutes and supernatants collected. 30 μg of whole cell extracts were treated with sample buffer and boiled at 95°C for 5 minutes. Samples were separated via SDS-PAGE gel electrophoresis (Biorad) and transferred to a nitrocellulose membrane with the Trans-Blot Turbo system (Biorad) per manufacturer instructions. Membranes were blocked with StartingBlock (Thermo Fisher) blocking buffer for 60 minutes at room temperature, followed by incubation with primary antibodies to detect PAX8 (Novus 32440, 1:1000 dilution), SOX17 (Abcam ab224637, 1:2,000), MECOM (Cell Signal C50E12, 1:1,000) or β-tubulin (Cell signaling D3U1W, 1:2,000). Primary antibody incubations were performed in blocking buffer overnight at 4°C. Samples were then washed with TBS-T three times for 10 minutes each and incubated in secondary antibody (1:10,000, Abcam ab6721 or ab6789) for 1 hour followed by three 10-minute TBS-T washes. Membranes were developed using Piece ECL Western Substrate (Thermo Scientific) following the manufacturer’s protocol.

#### RNA-sequencing and data analysis

OVCAR4 cells were transfected in triplicate. RNA and protein were harvested 72 hours-post transfection. Protein lysates were used to verify knockdown using western blotting, as described above. Cells were washed with cold PBS, collected by scraping and RNA extraction performed using the Nucleospin RNA Plus kit (Macherey-Nagel) per manufacturer’s protocol. Extracted RNA samples were used for poly-A non-stranded library preparation and 150 bp paired-end sequencing at 40million reads using the DNB-seq next generation sequencing platform (RNA-seq performed by BGI). Reads were filtered and aligned using STAR-2.5.1b (ref_genome_hg38_gencodev26) and a gene-level read count matrix generated using featureCounts (subread-1.6.3-source). Differential gene expression analyses were then performed using the R package DESeq2 (version 1.24.0). Differentially expressed genes were selected using an absolute log2 fold change ≥ 1 and adjusted p-value ≤ 0.01. Pathway analyses were performed using Metascape (metascape.org) (Zhou et al., 2019).

#### Proximity Ligation Assay

To perform the proximity ligation assay (PLA) we employed the Duolink Technology (DUO92101, Sigma-Aldrich). Kuramochi cells were grown for 24 hours on a 96-well imaging plate (0030741030, Eppendorf). Cells were fixed in 4% paraformaldehyde for 15 min, permeabilized with 0.25% Triton X-100 for 15 min, and blocked with 1% BSA in PBS containing 0.1% Tween-20 for 30 min. Primary antibodies against PAX8 (Novus, NBP2-29903, dilution 1:250), SOX17 (Cell Signaling, 81778S, dilution 1:250) and MECOM (Proteintech, 23201-1-AP, dilution 1:250) were incubated overnight at 4 °C. After three 5 min washes with TBST, PLA probes were incubated overnight at 4°C. Detection was performed using Duolink RED detection reagents as recommended by manufacturer. Samples were air dried and covered with Duolink mounting medium with DAPI and them imaged using a Nikon Eclipse Ti inverted microscope under ×40 magnification.

#### ChIP-seq of PAX8, SOX17, MECOM, and CTCF in Kuramochi cells

Kuramochi cells were grown to 80% confluence, cross-linked with 1% formaldehyde in PBS for 15 minutes, pelleted, and flash frozen. 100 million cells were used per ChIP with 10 μg of each antibody - PAX8 (Cell Signaling, catalogue number 59019, lot 1), SOX17 (R&D systems, catalogue number AF1924, lot KG0818071), MECOM (Cell Signaling, catalogue number 2593, lot 4). Sonications were performed with a qSonica microtip sonicator with 4 minutes total (30 sec ON, 1 min OFF), 18-21 watts. The supernatant of the sonicated lysates was incubated overnight at 4°C with the antibody and Invitrogen DynaI magnetic bead mix. After extensive washing, enriched chromatin was purified as follows - for PAX8, using a phenol:chloroform:isoamyl alcohol extraction (Boija et al., 2018), for SOX17 and MECOM, bead:antibody:chromatin complexes were resuspended in elution buffer, placed at 65°C for 45 minutes with intermittent vortexing and spun down. RNase A was added to the supernatant, and samples incubated at 65°C for 3.5 hours before a proteinase K digest at 42°C for 1 hour. DNA was then purified using PCR column purification. CTCF ChIP-Seq was performed as previously described (Schmidt et al., 2009) using 25 μg chromatin and 5 μg anti-CTCF antibody (Active Motif, catalogue number 61311) and chromatin sheared to 100-300 bp fragments using a Covaris E220 evolution Focused Ultrasonicator. ChIP libraries were constructed using the Kapa Hyper Library Preparation kit, quantified and sequenced on an Illumina NextSeq 500 sequencer. Reads were aligned to the hg19 version of the human reference genome using bowtie (Langmead et al., 2009) v1.2 with parameters -k 2 -m 2 –best and -l set to the read length. WIG files for display were made using MACS (Zhang et al., 2008) v1.4 with parameters -w -S –space=50 –nomodel –shiftsize=200. Regions statistically enriched in reads were identified using MACS v1.4 with corresponding input control and parameters -p 1e-9 –keep-dup=auto. Regions for the colocalization heatmap were constructed by collapsing regions enriched in PAX8, SOX17, and MECOM using bedtools merge (Quinlan and Hall, 2010) and creating 4kb regions centered on the center of the collapsed regions. Read coverage was quantified for heatmap analysis using bamToGFF (https://github.com/BradnerLab/pipeline) with parameters –m 100 -r using a mapped read bam with non-PCR duplicate reads created with samtools rmdup (Li et al., 2009). Heatmaps were ranked using read coverage quantified in 1 kb windows centered on the middle of each collapsed region using bamToGFF with parameters -m 1 -r.

#### THZ531, THZ1 and JQ1 Dose Response Curves

OVCAR4 cells were plated 96 well plates at 5,000 cells per well. After 24 hours, THZ531 (ApexBio), THZ1(Selleck Chemicals) and JQ1 (Tocris) or vehicle (DMSO) were added in 1:3 dilutions starting from 10000nM to 1.5nM in triplicate and incubated for 72 hours at 37°C. Cell numbers were quantified using the Promega Cell Titer Glo reagent. Signals were then normalized to the lowest dose, and IC50s were calculated with GraphPad Prism.

#### THZ531, THZ1 and JQ1 Drug Treatment for RNA

400,000 OVCAR4 cells were seeded in 60mm dishes 24 hours prior to experiment. Cells were treated with either low-dose THZ1 (50nM), high-dose THZ1 (250nM), low-dose THZ531 (100nM), high-dose THZ531 (500nM), JQ1 (500nM), or DMSO (500nM) for 6 hours. Plates were then washed with ice-cold PBS once followed by RNA extraction with Nucleospin RNA Mini Kit (Macherey-Nagel), followed by qPCR. Relative expression was measured normalized against DMSO control.

#### THZ531, THZ1 and JQ1 Global Transcriptome Data

RNA-seq data from THZ1-treated Kuramochi cells were obtained from GSE116282 (Zeng et al., 2018). Data were filtered to remove lowly transcripts (RPKM >1), to select protein coding genes and expression of the top 10% of genes, at 50 and 250 nM was plotted.

#### Quantitative PCR

Total RNA was converted to cDNA using random primers (Promega) and M-MLV Reverse Transcriptase RNase H (Promega) as per manufacturer’s instructions. cDNA was then amplified by QuantStudio 12K Flex Real-Time PCR system (Thermo Fisher) with Taqman Universal Master Mix with UNG (Applied Biosystems) along with the following probes: GAPDH, TUBB, and/or ACTB as housekeeping genes (Hs02786624_g1, Hs00742828_s1, and/or Hs01060665_g1) PAX8, SOX17, and MECOM (Hs00247586_m1,Hs00751752_s1, and Hs00602795_m1).

## Bibliography

Adler, E.K., Corona, R.I., Lee, J.M., Rodriguez-Malave, N., Mhawech-Fauceglia, P., Sowter, H., Hazelett, D.J., Lawrenson, K., and Gayther, S.A. (2017). The PAX8 cistrome in epithelial ovarian cancer. Oncotarget 8, 108316–108332.

Annala, M., Taavitsainen, S., Vandekerkhove, G., Bacon, J.V.W., Beja, K., Chi, K.N., Nykter, M., and Wyatt, A.W. (2018). Frequent mutation of the FOXA1 untranslated region in prostate cancer. Commun. Biol. 1, 122.

Baratta, M.G., Schinzel, A.C., Zwang, Y., Bandopadhayay, P., Bowman-Colin, C., Kutt, J., Curtis, J., Piao, H., Wong, L.C., Kung, A.L., et al. (2015). An in-tumor genetic screen reveals that the BET bromodomain protein, BRD4, is a potential therapeutic target in ovarian carcinoma. Proc Natl Acad Sci USA 112, 232–237.

Boija, A., Klein, I.A., Sabari, B.R., Dall’Agnese, A., Coffey, E.L., Zamudio, A.V., Li, C.H., Shrinivas, K., Manteiga, J.C., Hannett, N.M., et al. (2018). Transcription Factors Activate Genes through the Phase-Separation Capacity of Their Activation Domains. Cell 175, 1842–1855.e16.

Bradner, J.E., Hnisz, D., and Young, R.A. (2017). Transcriptional addiction in cancer. Cell 168, 629–643.

Brody, S.L., Yan, X.H., Wuerffel, M.K., Song, S.K., and Shapiro, S.D. (2000). Ciliogenesis and left-right axis defects in forkhead factor HFH-4-null mice. Am. J. Respir. Cell Mol. Biol. 23, 45–51.

Buchwalter, G., Hickey, M.M., Cromer, A., Selfors, L.M., Gunawardane, R.N., Frishman, J., Jeselsohn, R., Lim, E., Chi, D., Fu, X., et al. (2013). PDEF promotes luminal differentiation and acts as a survival factor for ER-positive breast cancer cells. Cancer Cell 23, 753–767.

Cancer Genome Atlas Research Network. Electronicaddress, and Cancer Genome Atlas Research Network (2017). Integrated genomic characterization of pancreatic ductal adenocarcinoma. Cancer Cell 32, 185–203.e13.

Cancer Genome Atlas Research Network. Electronicaddress, and Cancer Genome Atlas Research Network (2017). Comprehensive and integrated genomic characterization of adult soft tissue sarcomas. Cell 171, 950–965.e28.

Cancer Genome Atlas Research Network (2011). Integrated genomic analyses of ovarian carcinoma. Nature 474, 609–615.

Cancer Genome Atlas Research Network (2013). Comprehensive molecular characterization of clear cell renal cell carcinoma. Nature 499, 43–49.

Cancer Genome Atlas Research Network (2014). Comprehensive molecular characterization of urothelial bladder carcinoma. Nature 507, 315–322.

Cancer Genome Atlas Research Network, Ley, T.J., Miller, C., Ding, L., Raphael, B.J., Mungall, A.J., Robertson, A.G., Hoadley, K., Triche, T.J., Laird, P.W., et al. (2013). Genomic and epigenomic landscapes of adult de novo acute myeloid leukemia. N. Engl. J. Med. 368, 2059–2074.

Cancer Genome Atlas Research Network, Linehan, W.M., Spellman, P.T., Ricketts, C.J., Creighton, C.J., Fei, S.S., Davis, C., Wheeler, D.A., Murray, B.A., Schmidt, L., et al. (2016). Comprehensive Molecular Characterization of Papillary Renal-Cell Carcinoma. N. Engl. J. Med. 374, 135–145.

Cancer Genome Atlas Research Network, Analysis Working Group: Asan University, BC Cancer Agency, Brigham and Women’s Hospital, Broad Institute, Brown University, Case Western Reserve University, Dana-Farber Cancer Institute, Duke University, Greater Poland Cancer Centre, et al. (2017a). Integrated genomic characterization of oesophageal carcinoma. Nature 541, 169–175.

Cancer Genome Atlas Research Network, Albert Einstein College of Medicine, Analytical Biological Services, Barretos Cancer Hospital, Baylor College of Medicine, Beckman Research Institute of City of Hope, Buck Institute for Research on Aging, Canada’s Michael Smith Genome Sciences Centre, Harvard Medical School, Helen F. Graham Cancer Center &Research Institute at Christiana Care Health Services, et al. (2017b). Integrated genomic and molecular characterization of cervical cancer. Nature 543, 378–384.

Cerami, E., Gao, J., Dogrusoz, U., Gross, B.E., Sumer, S.O., Aksoy, B.A., Jacobsen, A., Byrne, C.J., Heuer, M.L., Larsson, E., et al. (2012). The cBio cancer genomics portal: an open platform for exploring multidimensional cancer genomics data. Cancer Discov. 2, 401–404.

Chapuy, B., McKeown, M.R., Lin, C.Y., Monti, S., Roemer, M.G.M., Qi, J., Rahl, P.B., Sun, H.H., Yeda, K.T., Doench, J.G., et al. (2013). Discovery and characterization of super-enhancer-associated dependencies in diffuse large B cell lymphoma. Cancer Cell 24, 777–790.

Cheng, X.-H., Black, M., Ustiyan, V., Le, T., Fulford, L., Sridharan, A., Medvedovic, M., Kalinichenko, V.V., Whitsett, J.A., and Kalin, T.V. (2014). SPDEF inhibits prostate carcinogenesis by disrupting a positive feedback loop in regulation of the Foxm1 oncogene. PLoS Genet. 10, e1004656.

Chen, L., Huang, M., Plummer, J., Pan, J., Jiang, Y.Y., Yang, Q., Silva, T.C., Gull, N., Chen, S., Ding, L.W., et al. (2019a). Master transcription factors form interconnected circuitry and orchestrate transcriptional networks in oesophageal adenocarcinoma. Gut.

Chen, Y., Xu, L., Mayakonda, A., Huang, M.-L., Kanojia, D., Tan, T.Z., Dakle, P., Lin, R.Y.-T., Ke, X.-Y., Said, J.W., et al. (2019b). Bromodomain and extraterminal proteins foster the core transcriptional regulatory programs and confer vulnerability in liposarcoma. Nat. Commun. 10, 1353.

Cherniack, A.D., Shen, H., Walter, V., Stewart, C., Murray, B.A., Bowlby, R., Hu, X., Ling, S., Soslow, R.A., Broaddus, R.R., et al. (2017). Integrated molecular characterization of uterine carcinosarcoma. Cancer Cell 31, 411–423.

Cheung, H.W., Cowley, G.S., Weir, B.A., Boehm, J.S., Rusin, S., Scott, J.A., East, A., Ali, L.D., Lizotte, P.H., Wong, T.C., et al. (2011). Systematic investigation of genetic vulnerabilities across cancer cell lines reveals lineage-specific dependencies in ovarian cancer. Proc Natl Acad Sci USA 108, 12372–12377.

Ciriello, G., Gatza, M.L., Beck, A.H., Wilkerson, M.D., Rhie, S.K., Pastore, A., Zhang, H., McLellan, M., Yau, C., Kandoth, C., et al. (2015). Comprehensive molecular portraits of invasive lobular breast cancer. Cell 163, 506–519.

Colaprico, A., Silva, T.C., Olsen, C., Garofano, L., Cava, C., Garolini, D., Sabedot, T.S., Malta, T.M., Pagnotta, S.M., Castiglioni, I., et al. (2016). TCGAbiolinks: an R/Bioconductor package for integrative analysis of TCGA data. Nucleic Acids Res. 44, e71.

Corona, R.I., Seo, J.-H., Lin, X., Hazelett, D.J., Reddy, J., Abassi, F., Lin, Y.G., Mhawech-Fauceglia, P.Y., Lester, J., Shah, S.P., et al. (2019). Non-coding Somatic Mutations Converge on the PAX8 Pathway in Epithelial Ovarian Cancer. BioRxiv.

Davis, C.F., Ricketts, C.J., Wang, M., Yang, L., Cherniack, A.D., Shen, H., Buhay, C., Kang, H., Kim, S.C., Fahey, C.C., et al. (2014). The somatic genomic landscape of chromophobe renal cell carcinoma. Cancer Cell 26, 319–330.

Delmore, J.E., Issa, G.C., Lemieux, M.E., Rahl, P.B., Shi, J., Jacobs, H.M., Kastritis, E., Gilpatrick, T., Paranal, R.M., Qi, J., et al. (2011). BET bromodomain inhibition as a therapeutic strategy to target c-Myc. Cell 146, 904–917.

Denny, S.K., Yang, D., Chuang, C.-H., Brady, J.J., Lim, J.S., Grüner, B.M., Chiou, S.-H., Schep, A.N., Baral, J., Hamard, C., et al. (2016). Nfib Promotes Metastasis through a Widespread Increase in Chromatin Accessibility. Cell 166, 328–342.

Domcke, S., Sinha, R., Levine, D.A., Sander, C., and Schultz, N. (2013). Evaluating cell lines as tumour models by comparison of genomic profiles. Nat. Commun. 4, 2126.

Durbin, A.D., Zimmerman, M.W., Dharia, N.V., Abraham, B.J., Iniguez, A.B., Weichert-Leahey, N., He, S., Krill-Burger, J.M., Root, D.E., Vazquez, F., et al. (2018). Selective gene dependencies in MYCN-amplified neuroblastoma include the core transcriptional regulatory circuitry. Nat. Genet. 50, 1240–1246.

D’Alessio, A.C., Fan, Z.P., Wert, K.J., Baranov, P., Cohen, M.A., Saini, J.S., Cohick, E., Charniga, C., Dadon, D., Hannett, N.M., et al. (2015). A Systematic Approach to Identify Candidate Transcription Factors that Control Cell Identity. Stem Cell Reports 5, 763–775.

Eliades, P., Abraham, B.J., Ji, Z., Miller, D.M., Christensen, C.L., Kwiatkowski, N., Kumar, R., Njauw, C.N., Taylor, M., Miao, B., et al. (2018). High MITF Expression Is Associated with Super-Enhancers and Suppressed by CDK7 Inhibition in Melanoma. J. Invest. Dermatol. 138, 1582–1590.

Elias, K.M., Emori, M.M., Westerling, T., Long, H., Budina-Kolomets, A., Li, F., MacDuffie, E., Davis, M.R., Holman, A., Lawney, B., et al. (2016). Epigenetic remodeling regulates transcriptional changes between ovarian cancer and benign precursors. JCI Insight 1.

Farshidfar, F., Zheng, S., Gingras, M.-C., Newton, Y., Shih, J., Robertson, A.G., Hinoue, T., Hoadley, K.A., Gibb, E.A., Roszik, J., et al. (2017). Integrative Genomic Analysis of Cholangiocarcinoma Identifies Distinct IDH-Mutant Molecular Profiles. Cell Rep. 19, 2878–2880.

Federation, A.J., Polaski, D.R., Ott, C.J., Fan, A., Lin, C.Y., and Bradner, J.E. (2018). Identification of candidate master transcription factors within enhancer-centric transcriptional regulatory networks. BioRxiv.

Fishbein, L., Leshchiner, I., Walter, V., Danilova, L., Robertson, A.G., Johnson, A.R., Lichtenberg, T.M., Murray, B.A., Ghayee, H.K., Else, T., et al. (2017). Comprehensive molecular characterization of pheochromocytoma and paraganglioma. Cancer Cell 31, 181–193.

Francavilla, C., Lupia, M., Tsafou, K., Villa, A., Kowalczyk, K., Rakownikow Jersie-Christensen, R., Bertalot, G., Confalonieri, S., Brunak, S., Jensen, L.J., et al. (2017). Phosphoproteomics of primary cells reveals druggable kinase signatures in ovarian cancer. Cell Rep. 18, 3242–3256.

Frank, R., Scheffler, M., Merkelbach-Bruse, S., Ihle, M.A., Kron, A., Rauer, M., Ueckeroth, F., König, K., Michels, S., Fischer, R., et al. (2018). Clinical and Pathological Characteristics of KEAP1- and NFE2L2-Mutated Non-Small Cell Lung Carcinoma (NSCLC). Clin. Cancer Res. 24, 3087–3096.

Frietze, S., O’Geen, H., Littlepage, L.E., Simion, C., Sweeney, C.A., Farnham, P.J., and Krig, S.R. (2014). Global analysis of ZNF217 chromatin occupancy in the breast cancer cell genome reveals an association with ERalpha. BMC Genomics 15, 520.

Gao, J., Aksoy, B.A., Dogrusoz, U., Dresdner, G., Gross, B., Sumer, S.O., Sun, Y., Jacobsen, A., Sinha, R., Larsson, E., et al. (2013). Integrative analysis of complex cancer genomics and clinical profiles using the cBioPortal. Sci. Signal. 6, pl1.

Grasso, C.S., Wu, Y.-M., Robinson, D.R., Cao, X., Dhanasekaran, S.M., Khan, A.P., Quist, M.J., Jing, X., Lonigro, R.J., Brenner, J.C., et al. (2012). The mutational landscape of lethal castration-resistant prostate cancer. Nature 487, 239–243.

Hmeljak, J., Sanchez-Vega, F., Hoadley, K.A., Shih, J., Stewart, C., Heiman, D., Tarpey, P., Danilova, L., Drill, E., Gibb, E.A., et al. (2018). Integrative molecular characterization of malignant pleural mesothelioma. Cancer Discov. 8, 1548–1565.

Hoadley, K.A., Yau, C., Hinoue, T., Wolf, D.M., Lazar, A.J., Drill, E., Shen, R., Taylor, A.M., Cherniack, A.D., Thorsson, V., et al. (2018). Cell-of-Origin Patterns Dominate the Molecular Classification of 10,000 Tumors from 33 Types of Cancer. Cell 173, 291–304.e6.

Hu, S., Marineau, J.J., Rajagopal, N., Hamman, K.B., Choi, Y.J., Schmidt, D.R., Ke, N., Johannessen, L., Bradley, M.J., Orlando, D.A., et al. (2019). Discovery and Characterization of SY-1365, a Selective, Covalent Inhibitor of CDK7. Cancer Res. 79, 3479–3491.

Jiang, X., Finucane, H.K., Schumacher, F.R., Schmit, S.L., Tyrer, J.P., Han, Y., Michailidou, K., Lesseur, C., Kuchenbaecker, K.B., Dennis, J., et al. (2019). Shared heritability and functional enrichment across six solid cancers. Nat. Commun. 10, 431.

Karst, A.M., Jones, P.M., Vena, N., Ligon, A.H., Liu, J.F., Hirsch, M.S., Etemadmoghadam, D., Bowtell, D.D.L., and Drapkin, R. (2014). Cyclin E1 deregulation occurs early in secretory cell transformation to promote formation of fallopian tube-derived high-grade serous ovarian cancers. Cancer Res. 74, 1141–1152.

Kar, S.P., Beesley, J., Amin Al Olama, A., Michailidou, K., Tyrer, J., Kote-Jarai, Zs., Lawrenson, K., Lindstrom, S., Ramus, S.J., Thompson, D.J., et al. (2016). Genome-Wide Meta-Analyses of Breast, Ovarian, and Prostate Cancer Association Studies Identify Multiple New Susceptibility Loci Shared by at Least Two Cancer Types. Cancer Discov. 6, 1052–1067.

Köbel, M., Kalloger, S.E., Boyd, N., McKinney, S., Mehl, E., Palmer, C., Leung, S., Bowen, N.J., Ionescu, D.N., Rajput, A., et al. (2008). Ovarian carcinoma subtypes are different diseases: implications for biomarker studies. PLoS Med. 5, e232.

Kuhn, E., Wang, T.-L., Doberstein, K., Bahadirli-Talbott, A., Ayhan, A., Sehdev, A.S., Drapkin, R., Kurman, R.J., and Shih, I.-M. (2016). CCNE1 amplification and centrosome number abnormality in serous tubal intraepithelial carcinoma: further evidence supporting its role as a precursor of ovarian high-grade serous carcinoma. Mod. Pathol. 29, 1254–1261.

Kwiatkowski, N., Zhang, T., Rahl, P.B., Abraham, B.J., Reddy, J., Ficarro, S.B., Dastur, A., Amzallag, A., Ramaswamy, S., Tesar, B., et al. (2014). Targeting transcription regulation in cancer with a covalent CDK7 inhibitor. Nature 511, 616–620.

Lambert, S.A., Jolma, A., Campitelli, L.F., Das, P.K., Yin, Y., Albu, M., Chen, X., Taipale, J., Hughes, T.R., and Weirauch, M.T. (2018). The human transcription factors. Cell 175, 598–599.

Langmead, B., Trapnell, C., Pop, M., and Salzberg, S.L. (2009). Ultrafast and memory-efficient alignment of short DNA sequences to the human genome. Genome Biol. 10, R25.

Lee, H.-S., Jang, H.-J., Shah, R., Yoon, D., Hamaji, M., Wald, O., Lee, J.-S., Sugarbaker, D.J., and Burt, B.M. (2017). Genomic Analysis of Thymic Epithelial Tumors Identifies Novel Subtypes Associated with Distinct Clinical Features. Clin. Cancer Res. 23, 4855–4864.

Lin, C.Y., Erkek, S., Tong, Y., Yin, L., Federation, A.J., Zapatka, M., Haldipur, P., Kawauchi, D., Risch, T., Warnatz, H.-J., et al. (2016). Active medulloblastoma enhancers reveal subgroup-specific cellular origins. Nature 530, 57–62.

Littlepage, L.E., Adler, A.S., Kouros-Mehr, H., Huang, G., Chou, J., Krig, S.R., Griffith, O.L., Korkola, J.E., Qu, K., Lawson, D.A., et al. (2012). The transcription factor ZNF217 is a prognostic biomarker and therapeutic target during breast cancer progression. Cancer Discov. 2, 638–651.

Li, H., Handsaker, B., Wysoker, A., Fennell, T., Ruan, J., Homer, N., Marth, G., Abecasis, G., Durbin, R., and 1000 Genome Project Data Processing Subgroup (2009). The Sequence Alignment/Map format and SAMtools. Bioinformatics 25, 2078–2079.

Lovén, J., Hoke, H.A., Lin, C.Y., Lau, A., Orlando, D.A., Vakoc, C.R., Bradner, J.E., Lee, T.I., and Young, R.A. (2013). Selective inhibition of tumor oncogenes by disruption of super-enhancers. Cell 153, 320–334.

Mansour, M.R., Abraham, B.J., Anders, L., Berezovskaya, A., Gutierrez, A., Durbin, A.D., Etchin, J., Lawton, L., Sallan, S.E., Silverman, L.B., et al. (2014). Oncogene regulation. An oncogenic super-enhancer formed through somatic mutation of a noncoding intergenic element. Science 346, 1373–1377.

Matsuyama, A., Hisaoka, M., Shimajiri, S., Hayashi, T., Imamura, T., Ishida, T., Fukunaga, M., Fukuhara, T., Minato, H., Nakajima, T., et al. (2006). Molecular detection of FUS-CREB3L2 fusion transcripts in low-grade fibromyxoid sarcoma using formalin-fixed, paraffin-embedded tissue specimens. Am. J. Surg. Pathol. 30, 1077–1084.

Meyers, R.M., Bryan, J.G., McFarland, J.M., Weir, B.A., Sizemore, A.E., Xu, H., Dharia, N.V., Montgomery, P.G., Cowley, G.S., Pantel, S., et al. (2017). Computational correction of copy number effect improves specificity of CRISPR-Cas9 essentiality screens in cancer cells. Nat. Genet. 49, 1779–1784.

Moon, H.-G., Hwang, K.-T., Kim, J.-A., Kim, H.S., Lee, M.-J., Jung, E.-M., Ko, E., Han, W., and Noh, D.-Y. (2011). NFIB is a potential target for estrogen receptor-negative breast cancers. Mol. Oncol. 5, 538–544.

Mounir, M., Lucchetta, M., Silva, T.C., Olsen, C., Bontempi, G., Chen, X., Noushmehr, H., Colaprico, A., and Papaleo, E. (2019). New functionalities in the TCGAbiolinks package for the study and integration of cancer data from GDC and GTEx. PLoS Comput. Biol. 15, e1006701.

Ott, C.J., Federation, A.J., Schwartz, L.S., Kasar, S., Klitgaard, J.L., Lenci, R., Li, Q., Lawlor, M., Fernandes, S.M., Souza, A., et al. (2018). Enhancer architecture and essential core regulatory circuitry of chronic lymphocytic leukemia. Cancer Cell 34, 982–995.e7.

Parker, S.C.J., Stitzel, M.L., Taylor, D.L., Orozco, J.M., Erdos, M.R., Akiyama, J.A., van Bueren, K.L., Chines, P.S., Narisu, N., NISC Comparative Sequencing Program, et al. (2013). Chromatin stretch enhancer states drive cell-specific gene regulation and harbor human disease risk variants. Proc Natl Acad Sci USA 110, 17921–17926.

Park, J.-M., Kim, M.-Y., Kim, T.-H., Min, D.-K., Yang, G.E., and Ahn, Y.-H. (2018). Prolactin regulatory element-binding (PREB) protein regulates hepatic glucose homeostasis. Biochim. Biophys. Acta Mol. Basis Dis. 1864, 2097–2107.

Patch, A.-M., Christie, E.L., Etemadmoghadam, D., Garsed, D.W., George, J., Fereday, S., Nones, K., Cowin, P., Alsop, K., Bailey, P.J., et al. (2015). Whole-genome characterization of chemoresistant ovarian cancer. Nature 521, 489–494.

Polyak, K. (2007). Breast cancer: origins and evolution. J. Clin. Invest. 117, 3155–3163.

Pomerantz, M.M., Li, F., Takeda, D.Y., Lenci, R., Chonkar, A., Chabot, M., Cejas, P., Vazquez, F., Cook, J., Shivdasani, R.A., et al. (2015). The androgen receptor cistrome is extensively reprogrammed in human prostate tumorigenesis. Nat. Genet. 47, 1346–1351.

Quinlan, A.R., and Hall, I.M. (2010). BEDTools: a flexible suite of utilities for comparing genomic features. Bioinformatics 26, 841–842.

Rauscher, B., Henkel, L., Heigwer, F., and Boutros, M. (2019). Lineage specific core-regulatory circuits determine gene essentiality in cancer cells. BioRxiv.

Robertson, A.G., Shih, J., Yau, C., Gibb, E.A., Oba, J., Mungall, K.L., Hess, J.M., Uzunangelov, V., Walter, V., Danilova, L., et al. (2017). Integrative analysis identifies four molecular and clinical subsets in uveal melanoma. Cancer Cell 32, 204–220.e15.

Robinson, J.L.L., and Carroll, J.S. (2012). FoxA1 is a key mediator of hormonal response in breast and prostate cancer. Front Endocrinol (Lausanne) 3, 68.

Robinson, J.L.L., Holmes, K.A., and Carroll, J.S. (2013). FOXA1 mutations in hormone-dependent cancers. Front. Oncol. 3, 20.

Saint-André, V., Federation, A.J., Lin, C.Y., Abraham, B.J., Reddy, J., Lee, T.I., Bradner, J.E., and Young, R.A. (2016). Models of human core transcriptional regulatory circuitries. Genome Res. 26, 385–396.

Sanda, T., Lawton, L.N., Barrasa, M.I., Fan, Z.P., Kohlhammer, H., Gutierrez, A., Ma, W., Tatarek, J., Ahn, Y., Kelliher, M.A., et al. (2012). Core transcriptional regulatory circuit controlled by the TAL1 complex in human T cell acute lymphoblastic leukemia. Cancer Cell 22, 209–221.

Santagata, S., Ligon, K.L., and Hornick, J.L. (2007). Embryonic stem cell transcription factor signatures in the diagnosis of primary and metastatic germ cell tumors. Am. J. Surg. Pathol. 31, 836–845.

Schmidt, D., Wilson, M.D., Spyrou, C., Brown, G.D., Hadfield, J., and Odom, D.T. (2009). ChIP-seq: using high-throughput sequencing to discover protein-DNA interactions. Methods 48, 240–248.

Shang, S., Yang, J., Jazaeri, A.A., Duval, A.J., Tufan, T., Lopes Fischer, N., Benamar, M., Guessous, F., Lee, I., Campbell, R.M., et al. (2019). Chemotherapy-induced distal enhancers drive transcriptional program to maintain the chemoresistant state in ovarian cancer. Cancer Res.

Shaoxian, T., Baohua, Y., Xiaoli, X., Yufan, C., Xiaoyu, T., Hongfen, L., Rui, B., Xiangjie, S., Ruohong, S., and Wentao, Y. (2017). Characterisation of GATA3 expression in invasive breast cancer: differences in histological subtypes and immunohistochemically defined molecular subtypes. J. Clin. Pathol. 70, 926–934.

Shen, H., Shih, J., Hollern, D.P., Wang, L., Bowlby, R., Tickoo, S.K., Thorsson, V., Mungall, A.J., Newton, Y., Hegde, A.M., et al. (2018). Integrated molecular characterization of testicular germ cell tumors. Cell Rep. 23, 3392–3406.

Song, L., Wang, X., and Feng, Z. (2018). Overexpression of FOXM1 as a target for malignant progression of esophageal squamous cell carcinoma. Oncol. Lett. 15, 5910–5914.

Takata, A., Takiguchi, S., Okada, K., Takahashi, T., Kurokawa, Y., Yamasaki, M., Miyata, H., Nakajima, K., Mori, M., and Doki, Y. (2014). Clinicopathological and prognostic significance of FOXM1 expression in esophageal squamous cell carcinoma. Anticancer Res. 34, 2427–2432.

Tsherniak, A., Vazquez, F., Montgomery, P.G., Weir, B.A., Kryukov, G., Cowley, G.S., Gill, S., Harrington, W.F., Pantel, S., Krill-Burger, J.M., et al. (2017). Defining a cancer dependency map. Cell 170, 564–576.e16.

Wang, Y., Zhang, T., Kwiatkowski, N., Abraham, B.J., Lee, T.I., Xie, S., Yuzugullu, H., Von, T., Li, H., Lin, Z., et al. (2015). CDK7-dependent transcriptional addiction in triple-negative breast cancer. Cell 163, 174–186.

Whyte, W.A., Orlando, D.A., Hnisz, D., Abraham, B.J., Lin, C.Y., Kagey, M.H., Rahl, P.B., Lee, T.I., and Young, R.A. (2013). Master transcription factors and mediator establish super-enhancers at key cell identity genes. Cell 153, 307–319.

Xiong, D., Pan, J., Yin, Y., Jiang, H., Szabo, E., Lubet, R.A., Wang, Y., and You, M. (2018). Novel mutational landscapes and expression signatures of lung squamous cell carcinoma. Oncotarget 9, 7424–7441.

Yuan, J., Jiang, Y.-Y., Mayakonda, A., Huang, M., Ding, L.-W., Lin, H., Yu, F., Lu, Y., Loh, T.K.S., Chow, M., et al. (2017). Super-Enhancers Promote Transcriptional Dysregulation in Nasopharyngeal Carcinoma. Cancer Res. 77, 6614–6626.

Zeng, M., Kwiatkowski, N.P., Zhang, T., Nabet, B., Xu, M., Liang, Y., Quan, C., Wang, J., Hao, M., Palakurthi, S., et al. (2018). Targeting MYC dependency in ovarian cancer through inhibition of CDK7 and CDK12/13. Elife 7.

Zhang, T., Kwiatkowski, N., Olson, C.M., Dixon-Clarke, S.E., Abraham, B.J., Greifenberg, A.K., Ficarro, S.B., Elkins, J.M., Liang, Y., Hannett, N.M., et al. (2016). Covalent targeting of remote cysteine residues to develop CDK12 and CDK13 inhibitors. Nat. Chem. Biol. 12, 876–884.

Zhang, Y., Liu, T., Meyer, C.A., Eeckhoute, J., Johnson, D.S., Bernstein, B.E., Nusbaum, C., Myers, R.M., Brown, M., Li, W., et al. (2008). Model-based analysis of ChIP-Seq (MACS). Genome Biol. 9, R137.

Zhang, Y., Federation, A.J., Kim, S., O’Keefe, J.P., Lun, M., Xiang, D., Brown, J.D., and Steinhauser, M.L. (2018). Targeting nuclear receptor NR4A1-dependent adipocyte progenitor quiescence promotes metabolic adaptation to obesity. J. Clin. Invest. 128, 4898–4911.

Zhao, N., Cao, J., Xu, L., Tang, Q., Dobrolecki, L.E., Lv, X., Talukdar, M., Lu, Y., Wang, X., Hu, D.Z., et al. (2018). Pharmacological targeting of MYC-regulated IRE1/XBP1 pathway suppresses MYC-driven breast cancer. J. Clin. Invest. 128, 1283–1299.

Zhou, Y., Zhou, B., Pache, L., Chang, M., Khodabakhshi, A.H., Tanaseichuk, O., Benner, C., and Chanda, S.K. (2019). Metascape provides a biologist-oriented resource for the analysis of systems-level datasets. Nat. Commun. 10, 1523.

